# Single-cell lineage tracing of human neuromesoderm organoids reveals TBX6-mediated posterior mesoderm fate diversification

**DOI:** 10.64898/2026.07.29.741448

**Authors:** Yuanxin Liao, Cheng Chen, Meisi Li, Chuwei Wang, Li Wang, Miao Zhu, Yao Yao, Guangdun Peng

**Author notes:** These authors contributed equally (Yuanxin Liao, Cheng Chen, Meisi Li).

## Abstract

Neuromesodermal progenitors (NMPs) drive vertebrate posterior axis elongation, but the lineage hierarchy of human NMP derivatives remains poorly defined. Here, we integrated a CRISPR/Cas9-based single-cell lineage tracing system with paired multiomics profiling—including scRNA-seq and scATAC-seq—in human pluripotent stem cell-derived neuromesodermal organoids (NMOs) to reconstruct high-resolution lineage relationships spanning 50 days of differentiation. Our data analysis reveal that intermediate mesoderm (IM) and paraxial mesoderm (PM) originate from a shared presomitic mesoderm (PSM) progenitor downstream of NMPs, whereas lateral plate mesoderm (LPM) segregates early from committed mesodermal progenitors. Notably, TBX6 ablation disrupts both IM and somite development, demonstrating that TBX6 functions as an upstream regulator of these two lineages. Mechanistically, TBX6 loss arrests PSM at the progenitor stage and abolishes activation of development programs required for IM maturation. We further delineate neural crest differentiation pathways from NMPs, identifying pre-bifurcation molecular signatures that predict neural crest fate. This work provides a clonal-resolution lineage map of human NMP differentiation and advances our understanding of human posterior trunk development and the etiology of TBX6-associated congenital disorders affecting both the spine and the kidney.

## INTRODUCTION

The vertebrate body plan is established progressively along the anteroposterior axis during embryonic development. A key driver of posterior embryonic elongation is a population of bipotent stem cells known as neuromesodermal progenitors (NMPs)^1–4^. First identified in mouse embryos, NMPs are characterized by co-expression of the neural marker Sox2 and the mesodermal marker TBXT (Brachyury) ^2,3,5^. These cells generate both spinal cord tissue and mesodermal derivatives^6–10^. The discovery of NMPs has revised traditional germ layer theory by demonstrating that posterior neural tissues originate from dual origins, highlighting a distinct mechanism for posterior axis specification^4,11,12^.

Lineage tracing studies in mouse models have established that NMPs generate not only paraxial mesoderm (PM) and spinal cord tissues but also trunk neural crest cells^6,13^. More recently, mouse studies showed that the nephric mesenchyme—the embryonic precursor of the kidney—develops from *Tbx6*-expressing NMP derivatives^14^, raising the question of whether NMPs also give rise to intermediate mesoderm (IM) and other mesodermal subtypes. Despite abundant data from mice, direct extrapolation to humans is constrained by substantial species-specific differences in developmental regulatory networks.

In humans, pathogenic variants in TBX6 are associated with both congenital scoliosis (somite defects) and congenital anomalies of the kidney and urinary tract (CAKUT, IM defects)^15–17^, suggesting that TBX6 may govern the development of both PM and IM. However, the underlying developmental mechanisms remain uncharacterized.

Recent technological advances in single-cell RNA sequencing (scRNA-seq) and spatial transcriptomics have generated high-resolution molecular atlases of mouse and human embryos, capturing extensive transcriptional heterogeneity within NMPs and their early mesodermal progeny^18–21^. Although single-cell and spatial atlases of human gastrulation have annotated diverse early mesodermal subsets^22,23^, these profiling datasets lack clonal resolution to trace the origins of posterior IM and lateral plate mesoderm (LPM) subtypes. Meanwhile, direct investigation of early human embryos is limited by ethical constraints, and existing trunk organoid models fail to fully recapitulate human NMP lineage diversification. Collectively, these gaps hinder our mechanistic understanding of how human NMPs diversify into distinct mesodermal subtypes, as well as the functional role of TBX6 in this fate-specification process.

To bridge this gap, we adapted a CRISPR/Cas9-based lineage recording system (Cas9-Tracer)^24–30^ for use in human neuromesodermal organoids (NMOs), enabling simultaneous capture of single-cell transcriptomes, chromatin accessibility landscapes, and heritable clonal barcode histories. We defined the roadmap of mesodermal fate diversification from NMPs and uncovered the essential role of TBX6 in directing PSM-to-IM transition, and establishes a platform for interrogating the developmental origins of congenital axial and renal disorders.

## RESULTS

### Establishment of a molecularly defined NMO system for human posterior neuromesodermal development

Multiple protocols have been developed to generate NMP-associated organoids, including trunk-like structures (TLSs), and retinoic acid (RA)– gastruloids^31–33^. However, substantial variability in cellular composition and lineage proportions persists across different induction strategies^34^, and these models do not capture the full spectrum of neural and mesodermal subtypes that arise during human trunk development.

To recapitulate the self-organization properties of trunk development, we established a complex neuromesodermal organoid model by adapting a previously reported protocol^31^. Briefly, hPSCs were first directed toward trunk progenitors via 2D culture with FGF2 and CHIR99021 for 3 days to induce NMP fate. These progenitors were then dissociated and cultured in 3D suspension to form NMOs (**Figure 1A**). Brightfield imaging captured continuous morphological growth and axial elongation of NMOs across day 2 to day 60 of 3D culture (**Figure 1B**). Time-course qPCR showed that the neural progenitor marker *SOX2* increased steadily over time. For mesodermal lineages, the PM marker *MESP2* was strongly upregulated. The expression of IM marker *WT1* increased rapidly starting at day 4. The LPM marker *PRRX1* was highly expressed as early as day 2 (**Figure S1A**). Immunofluorescence analysis of NMOs revealed early polarized and regionalized patterning, with distinct domains expressing SOX2 (neural progenitor), TBX6 (PM), PAX2 (IM), and PRRX1 (LPM), indicating spontaneous self-organization of the neural and three major mesodermal lineages (**Figures 1C and S1B**).

**Figure 1.**
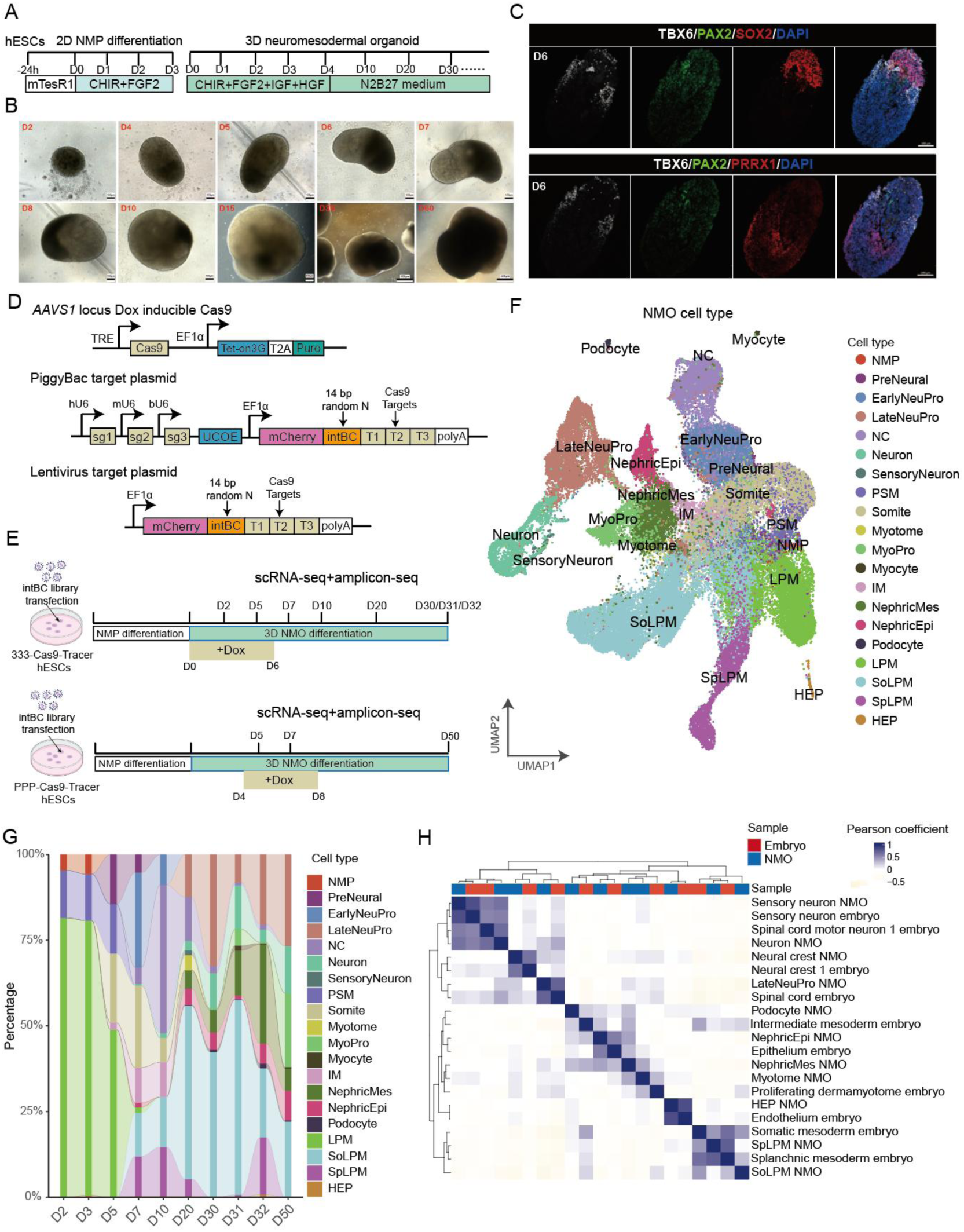
Single-cell lineage tracing reveals the developmental landscape of human neuromesodermal organoids. (A) Schematic of the NMO differentiation protocol, depicting the timeline of stepwise culture media conditions. 2D, two-dimensional; 3D, three-dimensional. (B) Brightfield images of NMOs at the indicated differentiation days, illustrating the morphological progression of organoid growth and self-organization during differentiation. Scale bar, 100-500 μm. (C) Immunofluorescence staining of two consecutive 20 μm thick serial sections from one day 6 (D6) NMO. Top section: TBX6 (paraxial mesoderm, green), PAX2 (intermediate mesoderm, red), SOX2 (neural lineage, grayscale), DAPI (blue); bottom adjacent section: TBX6 (paraxial mesoderm, green), PAX2 (intermediate mesoderm, red), PRRX1 (lateral plate mesoderm, red), DAPI (blue). Scale bar, 100 μm. (D) Schematic of the Cas9-Tracer system design. The system comprises three core genetic components stably integrated into human pluripotent stem cells: a doxycycline-inducible Cas9 expression construct integrated at the AAVS1 safe-harbor locus (top), and two distinct delivery vectors encoding the intBC reporter cassette (middle: piggybac transposon plasmid; bottom: lentiviral plasmid).For the PiggyBac target reporter plasmid: three independent sgRNA expression units (sg1/sg2/sg3) are each transcribed from species-specific U6 promoters (human hU6, mouse mU6, bovine bU6). Downstream, the EF1α promoter drives mCherry fluorescent reporter expression fused to a 14 bp random intBC, followed by three Cas9 cleavage target sites and a polyA terminator sequence. The lentiviral reporter backbone retains the identical mCherry-intBC-triple target cassette architecture but omits the three U6-sgRNA expression modules. (E) Experimental schematic of Cas9-Tracer lineage tracing across long-term NMO differentiation. Cas9-Tracer hESCs were first differentiated into NMPs via 2D culture, then NMPs were aggregated to generate 3D NMOs. Doxycycline was added to induce Cas9 expression, and samples were collected across 12 time points from D2 to D50. Paired scRNA-seq and targeted amplicon sequencing were processed to capture both transcriptomic profiles and lineage barcoding information. (F) UMAP of the integrated scRNA-seq dataset across all sampled developmental time points, colored by annotated cell types. NMP: neuromesodermal progenitors; PreNeural: preneural progenitors; EarlyNeuPro:early neural progenitors; LateNeuPro: late neural progenitors; NC: neural crest cells; PSM: presomitc mesoderm; MyoPro: Myocyte progenitor; IM: intermediate mesoderm; NephricMes: nephric mesenchyme; NephricEpi: nephric epithelium; LPM: lateral plate mesoderm; SoLPM: somatic lateral plate mesoderm; SpLPM: splanchnic lateral plate mesoderm. (G) Stacked bar plot showing the dynamic changes in cell type proportions across differentiation time points. All biological replicates collected at identical time points were combined and pooled prior to cell proportion quantification. (H) Pearson correlation heatmap comparing the transcriptome of NMO-derived cell types with primary human PCW4 cell types from published *in vivo* datasets.

To systematically characterize the cellular landscape of NMOs, we performed scRNA-seq on day 10 wild-type NMOs. Unsupervised clustering of scRNA-seq data uncovered a diverse spectrum of cell states, including neural progenitors, neural crest cells, and the three major mesodermal lineages: PM, IM, and LPM — each defined by canonical marker expression (**Figures S1C, S1D**). Immunofluorescence staining at day 10 further validated the spatial segregation of canonical lineage and trunk markers inside organoids (**Figures S1E–F**). Collectively, these findings confirm that NMOs possess the developmental potential to generate the major neural and mesodermal cell types of the human posterior trunk. This validates our organoid system as a reliable experimental platform for subsequent high-resolution lineage tracing analysis.

### Large-scale Cas9-Tracer lineage tracing in human NMOs resolves global posterior neuromesodermal lineage architecture

Classical lineage tracing assays in mouse embryos have verified that NMPs give rise to posterior neural tube tissues and paraxial mesoderm derivatives^6,35^. Nevertheless, the developmental potency of human NMPs remains largely uncharacterized. To define the global lineage hierarchy underlying human posterior neuromesodermal development, we performed unbiased, long-term clonal tracing across the entire time course of NMO differentiation using a Cas9-Tracer system adapted from Chan et al^36^ (**Figure 1D**). We employed two human pluripotent stem cell lines carrying the Cas9-Tracer cassette, each engineered with distinct sgRNA-target configurations: the 333 line (three mismatches) with moderate editing efficiency thus longer duration of recording, and the PPP line (no mismatch) with high editing efficiency (**Figures 1E, and Figure S2A, B**).

To capture lineage relationships across the full continuum of NMO maturation, we performed parallel, large-scale differentiation experiments using both Cas9-Tracer cell lines. For the PPP Cas9-Tracer cell line, Cas9 editing was induced by doxycycline at day 4 of 3D culture, whereas for the 333 Cas9-Tracer cell line, editing was induced at day 0 to enable earlier barcode accumulation. In total, we collected 12 samples across 10 developmental time points (days 2, 3, 5, 7, 10, 20, 30, 31, 32, and 50), covering the entire trajectory from early NMP emergence to terminal tissue maturation (**Figure 1E**). Brightfield and fluorescence imaging confirmed that Cas9-Tracer NMOs underwent morphologically normal elongation and self-organization, with consistent reporter expression throughout long-term culture (**Figure S2C**).

We then performed scRNA-seq and paired targeted amplicon sequencing on all 12 samples to capture both transcriptomic and lineage barcoding information. In total, we profiled around 69,500 single cells, with consistent cell yield across all samples (**Figures S3A-D**). Unsupervised integration of the transcriptome across all time points revealed that Cas9-Tracer NMOs generated the broad spectrum of neuromesodermal cell types, spanning NMPs, neural derivatives, and three major mesodermal lineages (**Figure 1F**) matching the cellular landscape of wild-type NMOs (Figure S1C, 1D).

We manually annotated 20 distinct cell types based on canonical marker expression, including neural populations of preneural progenitors (PreNeural), early/late neural progenitors (EarlyNeuPro/LateNeuPro), neural crest cells (NC), neurons, and sensory neurons and mesodermal populations of presomitic mesoderm (PSM), somites, myogenic derivatives, intermediate mesoderm (IM) and its nephric derivatives, and LPM subtypes **(Figure 1F)**. These populations were defined by canonical marker signatures: LPM (*FOXF1* and *PRRX1*), IM (*PAX2* and *PAX8*), PM (*TBX6* and *MEOX1*), neural progenitors (*SOX2*), sensory neurons (*POU4F1*), and neural crest (*SOX10*). Time-resolved analysis of cell-type proportions confirmed the expected lineage progression: early time points were dominated by NMPs and progressively lineage-restricted progenitors, which further were replaced by terminally differentiated cell types over time (**Figures 1G and S3G, H**). We further characterized posterior axial molecular signatures across D7, D20 and D50 NMOs by profiling *HOX* gene expression **(Figure S3F**). All neural and mesodermal cell subtypes displayed abundant expression of posterior *HOXB* and *HOXC* genes, with minimal activation of anterior marker *FOXG1*. This universal posterior *HOX* signature confirms that cell type within NMOs accurately recapitulating the axial molecular patterning of human posterior embryonic trunk tissues.

To validate the *in vivo* relevance of our NMO system, we compared the transcriptomes of our NMO-derived cell types at day 20 with published *in vivo* human embryonic cell datasets^37^. Pearson correlation analysis revealed high transcriptomic concordance between NMO-derived cell populations and their corresponding *in vivo* counterparts at PCW3 and PCW4 (**Figures 1H and S3G, H**), supporting the fidelity of our *in vitro* model. Together, these data establish a comprehensive single-cell atlas of human NMO differentiation and validate the system as a faithful model for dissecting NMP lineage diversification.

### Lineage bifurcation of NMP-derived neural progenitors into neural crest and neuron lineages

We first validated the lineage-tracing capacity of our Cas9-Tracer clonal recording platform across the full developmental time series of NMO differentiation. Across all sampled cell populations and developmental stages, 60% to 90% of single cell successfully recovered lineage barcodes (**Figure S3E**), demonstrating robust lineage recording efficiency. Each single cell recovered from all sampled time points carried an average of 3–6 unique integration barcodes (intBCs) (**Figure 2A**), with each intBC cassette harboring three independent Cas9 editing target sites to enable massive labelling capability. The frequency distribution of distinct intBC was relatively uniform at early differentiation stages. As organoid culture extended to late time points (D30–D50), clonal outgrowth drove skewed intBC abundance: select intBC clones acquired markedly higher cell proportions over prolonged organoid culture, a pattern consistent with stochastic clonal expansion to long-term 3D organoid growth (**Figures 2B** and **S2 D, E**). Calculation of clonal diversity indices based solely on intBC profiles confirmed sufficient barcode heterogeneity to resolve distinct ancestral progenitor populations across all developmental windows (**Figure S2G**).

**Figure 2.**
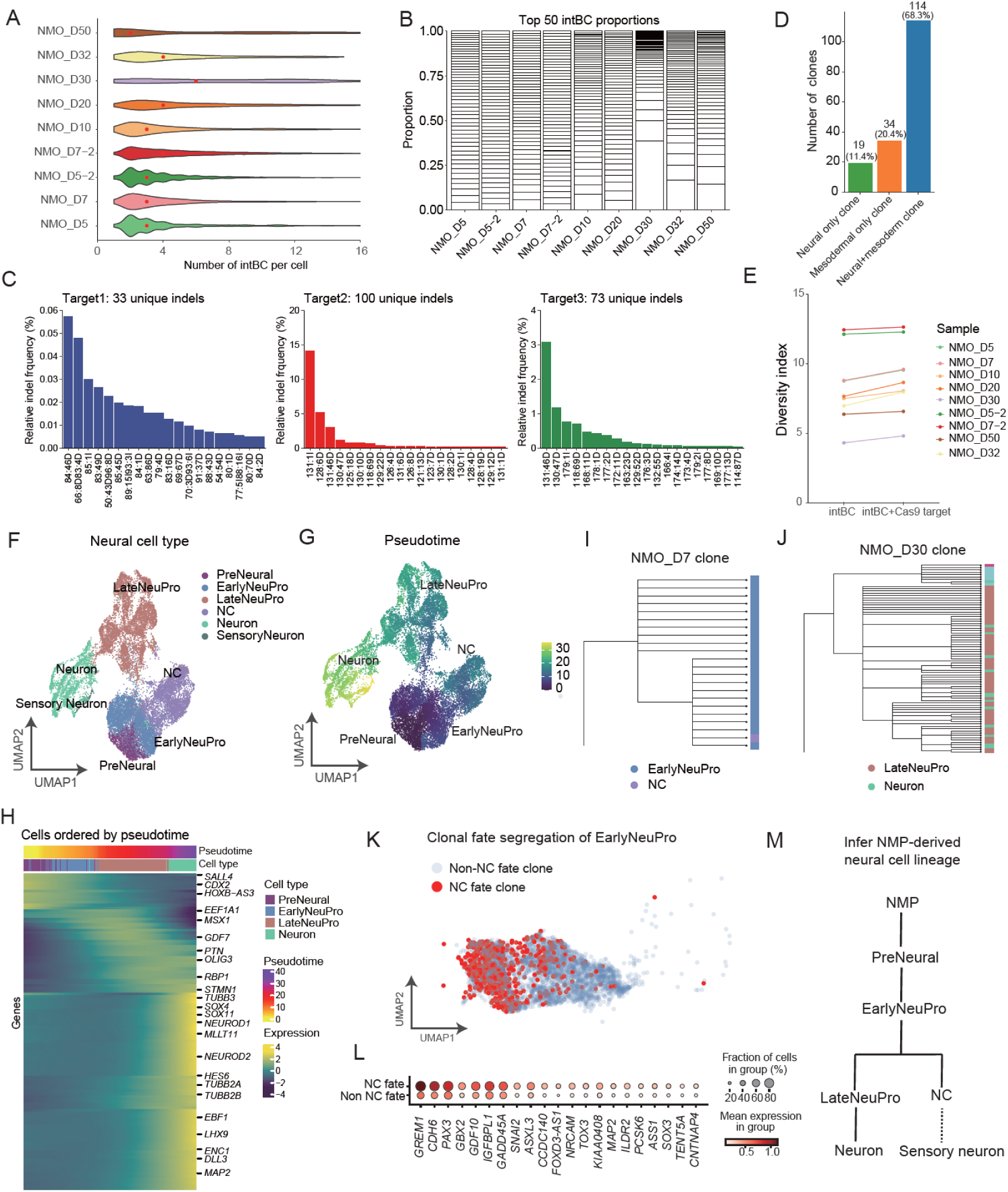
Clonal analysis resolves the early bifurcation of NMP-derived neural lineages. (A) Violin plots showing the distribution of intBC counts per cell across differentiation time points. (B) Stacked bar plot showing the relative proportions of the top 50 most abundant intBCs across differentiation time points from D5 to D50. (C) Bar plot showing the relative frequency of unique indels in the 333-Cas9-Tracer cell lines. Every indel label is formatted as alignment coordinate: indel size: indel class (I, insertions; D, deletions). Mutations were arranged in descending order of their read fractions, and the enumeration of unique indels was performed when the cumulative fraction first exceeded the 0.95 threshold. (D) Bar plot quantifying the fate composition of NMO clonal lineages, categorized into three classes: neural-only clones (green, n=19), mesodermal-only clones (blue, n=34), and mixed neural-mesodermal clones (orange, n=114). (E) Line plot showing the barcode diversity index across different differentiation time points. (F, G) UMAP of the re-clustered neural lineage cells, colored by cell type (F) and differentiation time point/sample (G). (H) Phylogenetic tree of a NMO day 7 clone, showing the lineage relationships of EarlyNeuPro and neural crest. (I) Phylogenetic tree of a NMO day 30 clone, showing the lineage relationships of LateNeuPro and neuron. (J) Heatmap showing the dynamic gene expression pattern along the pseudotime axis, with genes ordered by their expression dynamics, and cells ordered by pseudotime and cell type. (K) UMAP of the NMO D7 EarlyNeuPro cells, colored by whether the cells belong to a clone that contributes to NC cells. (L) Dot plot showing the top 20 differentially expressed genes between NC-committed and non-NC-committed EarlyNeuPro clones. Dot size indicates the fraction of cells expressing the gene, and color indicates the mean expression level. (M) Schematic model of the inferred NMP-derived neural lineage hierarchy. Solid lines represent direct lineage relationships supported by clonal barcode data. The dashed line indicates a previously reported developmental link between neural crest and sensory neurons, which was not directly captured by our clonal barcode.

Beyond static intBC labeling, doxycycline-induced Cas9 cleavage generates insertions and deletions (indels) to further enhance clonal discrimination. Peak editing efficiency across the three independent target loci reached approximately 50% within the sampled cell pool (**Figure S2F**). Sequencing of edited target amplicons recovered hundreds of unique indel signatures per target site (**Figures 2C and S2G, H**), which layered additional heritable sequence variation onto static intBC identifiers to resolve closely related descendant subclones originating from initial intBC-marked progenitor. Leveraging the combined combinatorial diversity of fixed intBC barcodes and progressive Cas9-generated indel scars, we reconstructed comprehensive clonal trees tracking lineage descendant throughout NMO development (**Figures S4A, B**). Consistent with the established bipotent nature of NMPs, clonal classification revealed that over two-thirds of recovered NMP-derived clones produced both neural and mesodermal progeny, with some clones restricted exclusively to neural or mesodermal fates **(Figure 2D)**.

Building on this global NMO lineage landscape, we next investigated neural fate decisions of human NMPs. Neuromesodermal progenitors give rise to posterior neural derivatives, including spinal motor neurons and neural crest cells^13,38^. Although NMP neural potency is well characterized in mice^39^, the precise lineage hierarchy and early regulatory mechanisms governing human NMP-derived neural differentiation remain poorly defined.

To address this, we isolated all neural lineage cells from our scRNA-seq dataset (**Figures 2F and S5A-F**). Pseudotime trajectory analysis resolved the lineage relationships between neuronal states, revealing two distinct differentiation paths originating from the EarlyNeuPro bifurcation node. One path directly transitioned from EarlyNeuPro to NC cells (**Figures S5D-F**), while the alternative path first progressed through LateNeuPro progenitors before terminally differentiating into functional neurons (**Figures 2G** and **S5A-C**). We further profiled dynamic gene expression along the pseudotime axis and observed sequential activation of neuron-specific regulatory genes during maturation (**Figure 2H**). Hierarchical clustering of NMO cell types at day 7 and day 30 further supported the lineage proximity between EarlyNeuPro and NC cells, as well as between LateNeuPro and neurons (**Figures 2I, J**).

To uncover molecular signatures that mark cell fate bias prior to neural lineage bifurcation, we stratified EarlyNeuPro cells based on their clonal origins using barcode information. We divided these progenitors into two groups: clones that ultimately generated neural crest (NC clones) and clones restricted exclusively to the neuronal lineage (non-NC clones) (**Figure 2K**). Differential gene expression analysis revealed that these two groups already exhibited significant transcriptional differences at the EarlyNeuPro stage, before clear lineage separation. Specifically, NC clones showed high expression of known NC regulators including *GREM1* and *PAX3*, consistent with their well-established roles in NC specification^40–43^. We also identified several novel pre-bifurcation genes, including *GADD45A* and *GBX2*, which were significantly upregulated in NC-committed EarlyNeuPro, suggesting that they may function as potential regulators of NC fate specification (**Figure 2L**). Notably, the top differentially expressed genes, including *GREM1*, *CHD6*, *PAX3*, and *GADD45A*, were highly expressed in subsequent neural crest cells, providing additional evidence supporting their potential role in early NC fate specification (**Figure S5H**).

Together, these results demonstrate that our NMO system recapitulates the full neural differentiation program of human NMPs. Our integrated clonal analysis confirms the conserved lineage architecture between human and mouse posterior neural development, and also identifies pre-bifurcation transcriptional signatures that may predict NC fate.

### State-fate mapping reveals NMP plasticity and the origin of posterior LPM

Following the characterization of neural lineage hierarchies, we next focused on resolving the lineage origins of three major posterior mesodermal subtypes: PM, IM and LPM. We first analyzed the clonal fates of mesoderm-restricted clones (≥10 cells per clone). Of these 34 purely mesodermal-restricted clones, 11 clones (32.4%) were strictly LPM-restricted, producing only LPM descendants with no PM, IM or neural progeny. By contrast, we recovered just a single PM-restricted clone, and zero clones that generated solely IM (**Figure 3A**). UMAP visualization further confirmed that descendants of representative LPM-restricted clones were spatially segregated from PM and IM populations (**Figures 3B–C**).

**Figure 3.**
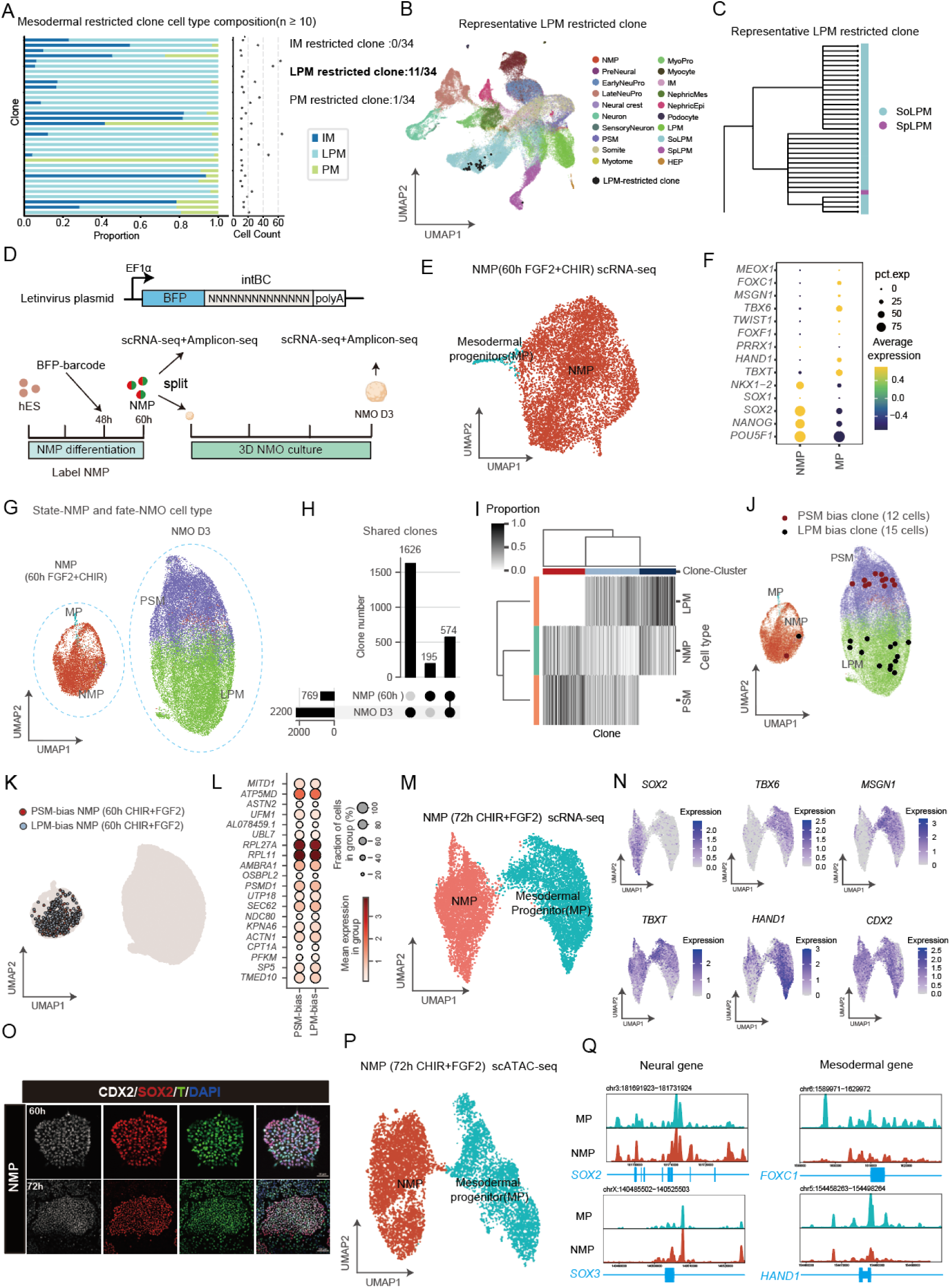
State-fate mapping reveals NMP plasticity and the origin of posterior lateral plate mesoderm. (A) Heatmap showing the clonal composition of mesodermal-restricted clones (n ≥ 10 cells per clone), with each row representing an individual clone and columns representing the proportion of cells in each mesodermal subtype (IM, LPM, PM). Clones are grouped by their exclusive fate output: IM-restricted, LPM-restricted, or PM-restricted. A dot plot on the right indicates the total cell count per clone. (B) Cells from a representative LPM-restricted clone are highlighted in red in the integration UMAP. (C) Phylogenetic tree of LPM-restricted clones, showing the hierarchical relationship between SoLPM and SpLPM derivatives. (D) Schematic of the state-fate mapping experiment. (E) UMAP of scRNA-seq data from 60 h (hours) NMP, colored by cell type, showing the NMP and MP populations. (F) Dot plot showing differentially expressed genes between MP and NMP populations at day 3. (G) UMAP of scRNA-seq data from the initial 60 h (hours) NMP (left) and final day 3 NMO derivatives (right), colored by annotated cell type. (H) Bar plot showing the number of shared barcoded clones recovered between the initial NMP and final time points NMO D3. (I) Heatmap showing the fate composition of individual NMP-derived clones, ordered by their fate bias, with rows representing clones and columns representing cell types. (J) UMAP highlighting two representative NMP-derived clones: the PSM-bias clone (red) and the LPM-bias clone (black), showing the distribution of their descendant cells in NMO D3. (K) UMAP of the NMPs, with NMP clones colored by their inferred fate bias (based on their descendant cell composition). (L) Dot plot showing the differential gene expression analysis between PSM-bias and LPM-bias NMPs at day 3. (M) UMAP of scRNA-seq data from 72 h (hours) NMPs, showing two distinct populations: NMPs (left) and mesodermal progenitors (MPs, right). (N) UMAP feature plots showing the expression of canonical marker genes (*SOX2, TBX6, MSGN1, TBXT, HAND1, CDX2*) across the NMP and MP populations. (O) UMAP of scATAC-seq data from 72h (hours) NMPs, identifying the same NMP and MP populations observed in the scRNA-seq data. (P) Genome browser tracks showing chromatin accessibility at neural gene loci and mesodermal gene loci in the NMP and MP populations. (Q) Immunofluorescence staining of NMP at 60 h (hours) and 72 h (hours), confirming the generation of NMPs via co-expression of the TBXT (Brachyury, red), neural marker SOX2 (green), and posterior trunk marker CDX2 (gray), with DAPI nuclear counterstaining (blue). Scale bar, 100 μm.

The concurrent emergence of LPM, PM and IM in NMOs raised a key question regarding the developmental origin of LPM: whether it derives directly from canonical NMPs or arises from specialized mesodermal progenitors specified prior to NMP formation. To address this issue, we performed high-resolution barcode-based state-fate mapping. We transduced human embryonic stem cell (hESC)-derived NMPs with a BFP-tagged intBC lentiviral library at 48 hours of differentiation, when cultures were highly enriched for nascent NMPs. After 12 hours, cells were divided into two parallel groups: one group was immediately harvested for scRNA-seq and amplicon sequencing, to record the initial transcriptional profiles and unique barcode signatures of individual NMPs; the other group was assembled into 3D NMOs and cultured for an additional three days. Subsequent scRNA-seq and barcode sequencing were performed to track terminal cell fate outcomes. State-fate correlation of NMPs was reconstructed by cross-matching barcode information obtained from the initial progenitor state and mature organoid derivatives (**Figure 3D**).

scRNA-seq analysis of cells collected at 60 hours confirmed barcoded cells were predominantly *SOX2* and *TBXT* double-positive NMPs, with a minor subset of committed mesodermal progenitors (MPs) (**Figures 3E, F**). We then matched barcodes between the initial NMP progenitors and the barcoded day 3 NMO derivatives to reconstruct the fate of each clone. A total of 574 overlapping clonal lineages were recovered, offering reliable statistical support for analyzing clonal fate preferences (**Figures 3G, H**).

Notably, NMP-derived clones gave rise not only to PSM, as previously reported^44^, but also to LPM cells, demonstrating that NMPs possess the developmental potential to generate posterior LPM in this model (**Figure 3I**). We further classified NMP-derived clones based on their fate composition, identifying two distinct groups: clones with a strong bias toward PSM, and clones with a bias toward LPM (**Figure 3J**). To examine whether such fate preference is predetermined at the transcriptional level, we performed differential gene expression analysis between the two clone groups. No significant transcriptional discrepancies were observed between PSM-biased and LPM-biased NMP clones (**Figures 3K, L**). This indicates that early NMPs remain transcriptionally uncommitted, and their terminal fate choices are likely regulated by extrinsic microenvironmental cues or epigenetic modifications rather than pre-established intrinsic transcriptional programs at this stage.

To identify the intermediate cell population that mediates LPM segregation downstream of uncommitted NMPs, we extended our single-cell profiling to the 72h differentiation time point, a window when mesodermal commitment becomes molecularly apparent. Unsupervised clustering of 72 h scRNA-seq data resolved two distinct cell states: the original bipotent NMP compartment (*SOX2⁺TBXT⁺*) and the fate-restricted mesodermal progenitor (MP) **(Figure 3M)**. A defining transcriptional signature of MPs is marked suppression of the neural master regulator SOX2, while these cells maintain high expression of core mesodermal transcription factors including TBXT, TBX6 and HAND1 **(Figure 3N)**. This transcriptional shift suggests MPs emerge as committed derivatives of bipotent NMPs: upon initiating mesodermal differentiation, NMPs downregulate SOX2, lose their dual neural-mesodermal potency, and lock into exclusive mesodermal differentiation programs.

We validated this temporal and spatial separation of NMP and MP states via whole-mount immunofluorescence of adherent NMP induction cultures, comparing the 60 h and 72 h time points **(Figure 3O)**. At 60 h, the monolayer culture consisted almost entirely of uniformly distributed *SOX2⁺TBXT⁺* dual-positive NMPs, with no distinct mesodermal subpopulation visible. By 72 h of differentiation, a discrete peripheral ring of *TBXT*-single-positive, –negative cells emerged surrounding a central core of bipotent NMPs; this outer spatially restricted population corresponds directly to the MP cluster captured in our 72h scRNA-seq.

To further interrogate the epigenetic regulatory differences that stabilize NMP and MP cell identities, we integrated parallel scATAC-seq profiling on 72 h NMP differentiation. scATAC-seq results further revealed selective chromatin opening at mesodermal gene loci in MPs, supporting their mesodermal commitment (**Figure 3P, Q**). Collectively, these findings suggest that the early-segregating LPM lineage primarily originates from committed MPs, which explains the widespread existence of LPM-only clones in our system.

Together, these results demonstrate that posterior LPM can be derived from NMPs *in vitro*, indicating that early NMPs retain broad developmental plasticity prior to fate restriction. Consistent with our observations, a recent preprint similarly reported that mouse NMPs harbor latent LPM potential^45^.

### Clonal analysis resolves the hierarchy of NMP-derived mesodermal lineage bifurcation

After defining the early segregation of LPM from NMPs, we next dissected the clonal lineage architecture governing differentiation of the remaining two major posterior mesodermal subtypes: PM (somite precursors) and IM (the embryonic precursor of kidney and urogenital tissues). The precise developmental cascade linking human NMPs to IM lineages remains poorly resolved, in contrast to well-characterized PM ontogeny in murine models^46^.

To resolve mesodermal lineage relationships, we reconstructed cell-type phylogenetic trees based on shared heritable Cas9 barcodes, which quantify clonal relatedness between terminal cell populations. Hierarchical clustering of day 7 NMO cell types demonstrated that IM and somite cells form a tightly clustered monophyletic clade, indicating they derive from a common immediate progenitor (**Figure 4A**). By contrast, somatic and splanchnic LPM populations split at an earlier ancestral node, consistent with their early fate segregation upstream of the IM-PM bifurcation event. A circular phylogenetic tree built from day 32 clonal barcodes further validated this topology: all LPM-associated clonal branches diverge at the root of the tree, while IM and somite subclusters arise from late-stage shared progenitors (**Figure 4B**). Consistent with this phylogenetic signal, we recovered multiple individual clones that simultaneously gave rise to both IM and somite cells at day 5 of differentiation, providing direct clonal evidence for their shared developmental origin (**Figure 4C**).

**Figure 4.**
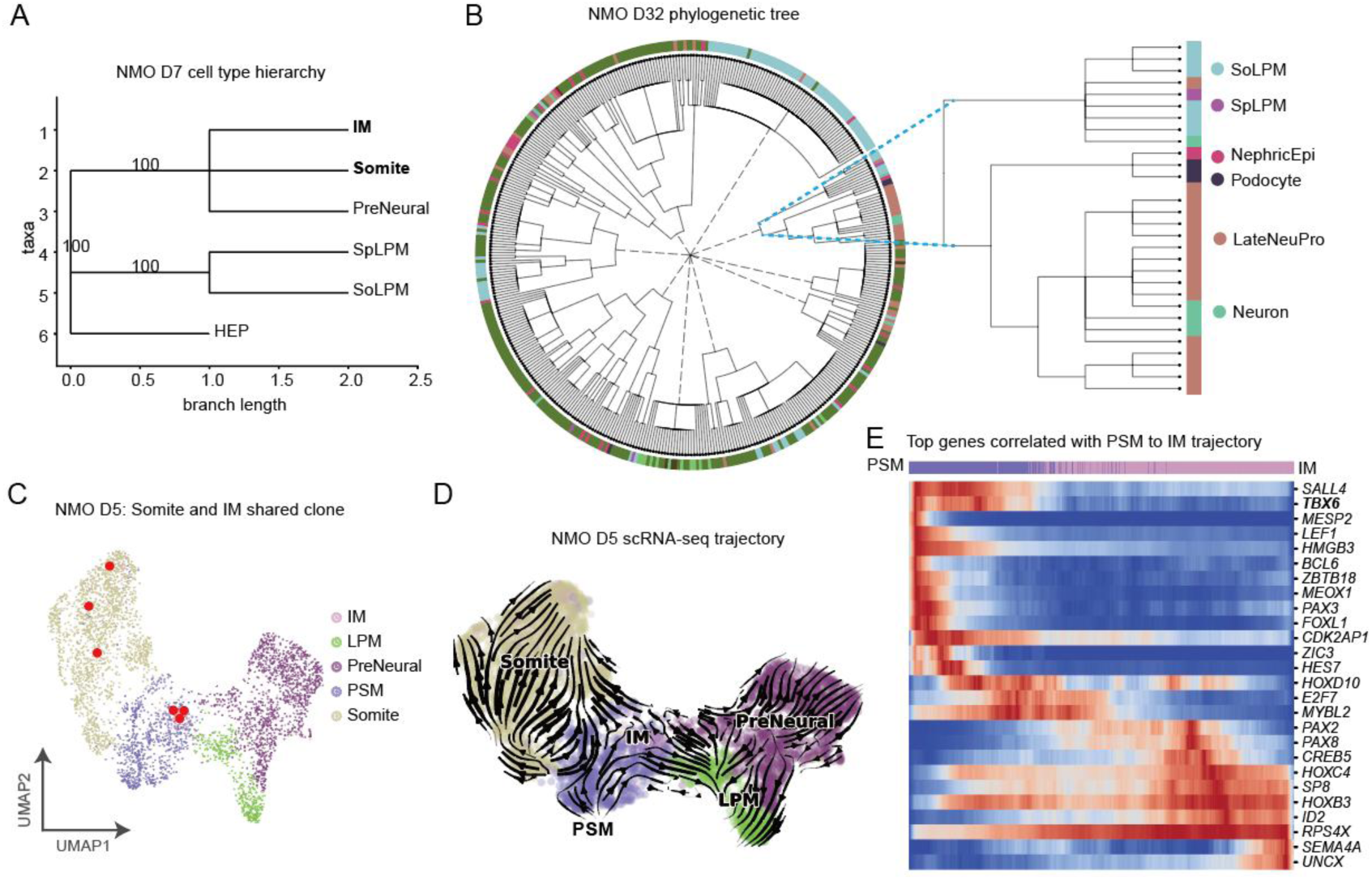
Clonal analysis resolves the hierarchy of NMP-derived mesodermal lineage bifurcation. (A) Hierarchical clustering tree of NMO cell types at day 7 (D7), based on lineage barcode. (B) Circular phylogenetic tree reconstructed from clonal barcodes at NMO day 32 (D32), showing the lineage relationships between terminal cell types. Each outer ring is colored by the terminal cell type, and dashed lines connect cells derived from different clones. (C) UMAP of NMO D5 scRNA-seq, colored by cell type. Cells from a representative IM and somite co-contributing clone are highlighted in red. (D) Pseudotime trajectory analysis of day 5 NMO cells. (E) Heatmap showing the top variable genes correlated with the PSM to IM trajectory, ordered by pseudotime.

To capture the continuous transcriptional trajectory preceding IM-PM fate split, we performed pseudotime ordering on NMO day 5 scRNA-seq data. The trajectory initiates from the PSM marked by canonical markers *TBX6* and *MSGN1*. From this PSM progenitor pool, two distinct mutually exclusive differentiation trajectories diverge: one transcriptional branch progresses toward intermediate mesoderm (IM), marked by gradual upregulation of IM signature genes including PAX2 and PAX8, and a second separate trajectory gives rise to somite (paraxial mesoderm). (**Figure 4D**). Differential gene ranking across the PSM-to-IM transition trajectory identified *TBX6* among the most dynamically enriched transcripts within the PSM compartment (**Figure 4E**).

### Loss of TBX6 causes presomitic mesoderm delay and impairs intermediate mesoderm maturation

Previous studies in mouse models have established that TBX6 is a critical regulator of paraxial mesoderm formation: loss of TBX6 leads to severe defects in somite formation, with presomitic cells aberrantly converting into neural derivatives^47–49^. However, TBX6 function in IM development remains less characterized in human systems. Based on these observations, we hypothesized that TBX6 is essential not only for somite formation but also for proper IM specification and maturation (**Figure 5A**).

**Figure 5.**
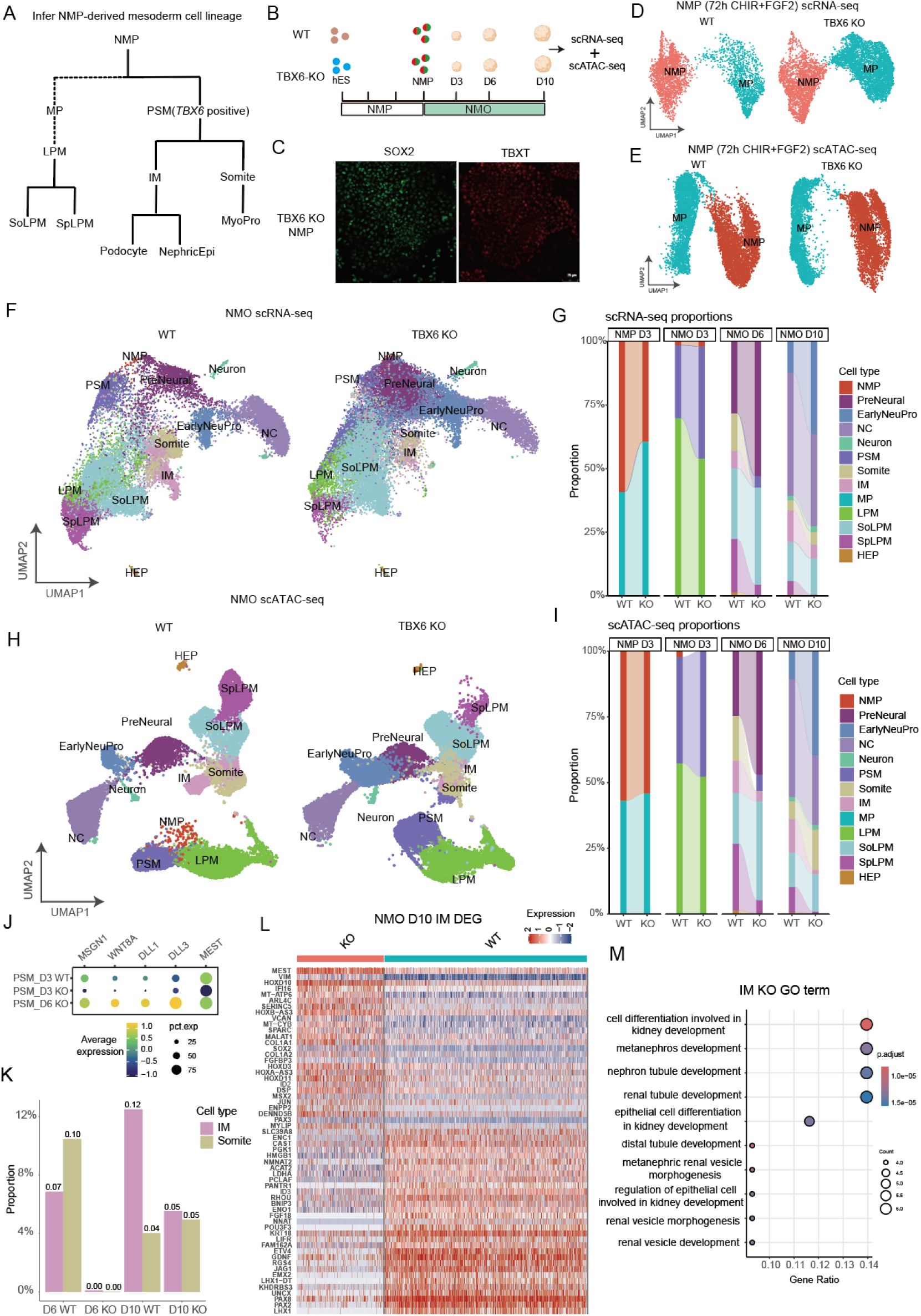
TBX6 Loss disrupts PSM progression and impairs intermediate mesoderm development. (A) Schematic model of the inferred NMP-derived mesodermal lineage hierarchy. (B) Schematic of the TBX6 KO experiment. Wild-type and TBX6-KO hESCs were differentiated into NMOs, with scRNA-seq and scATAC-seq collected at 72 h (hours) NMP and day 3, 6, 10 of NMO differentiation. (C) Immunofluorescence staining of 72 h (hours) NMP for SOX2 and TBXT, confirming the presence of SOX2⁺TBXT⁺ NMPs in TBX6-KO conditions. (D) UMAP of 72 h (hours) NMP scRNA-seq, colored by cell type. (E) UMAP of 72 h (hours) NMP scATAC-seq, colored by cell type. (F) UMAP of integrated scRNA-seq data from wild-type and TBX6-KO NMOs at days 3, 6, and 10, colored by cell type. (G) Stacked bar plot showing the proportion of each annotated cell type across differentiation time points (D3, D6, D10) in wild-type and TBX6-KO NMOs, based on scRNA-seq data. (H) UMAP of integrated scATAC-seq data from wild-type and TBX6-KO NMOs at days 3, 6, and 10, colored by cell type. (I) Stacked bar plot showing the proportion of each annotated cell type across differentiation time points (D3, D6, D10) in wild-type and TBX6-KO NMOs, based on scATAC-seq data. (J) Dot plot showing the average expression of cells expressing key PSM markers (*MEST, DLL3, DLL1, WNT8A, MSGN1*) in wild-type and TBX6-KO NMOs at day 3 and 6. (K) Bar plot showing the proportion of IM and somite cells in wild-type and TBX6-KO NMOs at day 6 and day 10, confirming the early loss of both lineages and the partial recovery of somite cells at day 10. (L) Heatmap showing differentially expressed genes (DEGs) of IM from day 10 wild-type and TBX6 KO NMOs. (M) Dot plot showing the gene ontology enrichment analysis of the downregulated genes in TBX6-KO IM cells, with terms related to kidney development highlighted.

To test this hypothesis, we generated a TBX6 homozygous knockout (KO) human pluripotent stem cell (hPSC) line via CRISPR/Cas9 editing and confirmed complete TBX6 protein loss by immunofluorescence (**Figures S6A, B**). We next examined whether TBX6 deficiency affects the induction of NMP. We found that TBX6-KO cells retained normal NMP induction capacity: at 72 h of differentiation, KO cells expressed SOX2^+^ and TBXT^+^ at levels comparable to wild-type (**Figure 5C**). Besides, scRNA-seq and scATAC-seq analysis confirmed the presence of *SOX2*^+^/*TBXT*^+^ NMPs (**Figures 5D, E and S6C, D**). Although scRNA-seq and scATAC-seq profiling of day 3 NMP revealed that WT and TBX6-KO NMP and MP formed transcriptionally and epigenetically distinct clusters with subtle differences (**Figures S6E, F**), the core NMP regulatory program remained intact, confirming that TBX6 is dispensable for NMP induction.

To dissect how TBX6 deletion reshapes human trunk mesodermal differentiation downstream of intact NMP specification, we subjected WT and TBX6-KO hPSCs to neuromesodermal organoid culture, harvesting matched samples at days 3, 6 and 10 to capture dynamic developmental defects (**Figure 5B**). We employed paired scRNA-seq and scATAC-seq to profile transcriptional and epigenomic cell-state changes in parallel, enabling orthogonal validation of lineage defects at two independent molecular layers. The comprehensive quality control for all scATAC-seq datasets; violin plots of TSS enrichment, fragment size distribution and normalized insertion profiles confirmed uniform high-quality chromatin accessibility profiling across all WT and KO time points (**Figure S8A–C**). Integrated UMAP embedding of scATAC-seq data recovered all canonical neuromesodermal cell subtypes in both genotypes, matching the cell-type annotation framework defined by our scRNA-seq atlas (**Figures S8D–E**).

Integrated UMAP visualization of cross-timepoint scRNA-seq datasets captured the full spectrum of human posterior trunk cell states in both WT and TBX6-KO NMOs, including neural progenitors, neural crest cells, PSM, IM, somites and LPM subtypes (**Figure 5F**). Despite recovery of all core cell populations, quantitative cell-type proportion tracking across days 3, 6 and 10 revealed dramatic, stage-dependent remodeling of mesodermal lineage differentiation upon TBX6 loss (**Figure 5G**). Parallel scATAC-seq UMAP embedding recapitulated this global cell composition shift: all major neural and mesodermal clusters were present in KO NMOs, yet the relative fraction of PSM, IM and somite populations was drastically altered relative to WT (**Figures 5H, I**). Log-fold change quantification of cell-type abundance shifts across differentiation stages further visualized progressive mesodermal depletion and neural lineage expansion in KO samples (**Figures S7C and S8F**).

Focused analysis of the PSM compartment—the shared progenitor pool for IM and somite lineages—identified early molecular defects initiating at day 3 of NMOs: robust downregulation of core PSM markers *MSGN1*, *DLX3* and *WNT8A* in day 3 KO PSM populations (**Figure 5J**). By day 6 of differentiation, WT PSM progenitors had fully exited the presomitic state and progressed to downstream IM and somite subtypes, rendering PSM nearly undetectable in WT NMOs. In contrast, PSM progenitors persisted in day 6 TBX6-KO NMOs, demonstrating that loss of TBX6 blocks PSM exit from the uncommitted progenitor state and prevents progression to terminal mesodermal fates.

Consistent with this arrest, TBX6-KO NMOs at day 6 showed near-complete loss of IM and somite populations, compared to 7% IM and 10% somite in WT controls (**Figure 5K**). Consistent results were observed in our scATAC-seq dataset, where the proportion of IM and somite cells was similarly depleted in KO NMO at day 6 (**Figure 5K**). We also observed a concomitant expansion of preneural and early neural progenitor populations in TBX6-KO NMO, a phenotype consistent with previous findings in mouse models^20,50^.

Partial somite recovery was observed by day 10, likely reflecting slow, TBX6-independent differentiation of arrested PSM cells (**Figure 5K**). In contrast, IM remained severely depleted at this later time point: only 5% of TBX6-KO NMOs were IM cells, versus 12% in WT. (**Figure 5K**). Moreover, we found that IM generated in KO NMOs exhibited distinct transcriptomic state: PCA analysis confirmed that KO IM formed a separate cluster from WT IM, with extensive changes in gene expression (**Figure S7F**). Differential gene expression analysis revealed that key IM and kidney progenitor genes, including *WT1*, *GDNF*, and *PAX8*, were substantially downregulated in TBX6-KO IM (**Figure 5L**). Gene Ontology (GO) enrichment analysis of the downregulated genes revealed strong enrichment for terms related to kidney and metanephros development, including nephric vesicle morphogenesis and kidney epithelium differentiation (**Figure 5M**).

To further assess the impact of TBX6 loss on lineage commitment, we compared the transcriptomic differences between somite and IM cells in WT and KO NMOs at day 10. In WT NMOs, these two lineages exhibited highly distinct transcriptional profiles, with numerous differentially expressed genes defining their unique lineage identities (**Figure S7F**). However, in TBX6-KO NMO, this lineage-specific transcriptional separation was severely attenuated: somite and IM had fewer differentially expressed genes, with key lineage-specific markers that distinguish IM from somite failing to be properly upregulated (**Figure S7G**).

Collectively, these data demonstrate that TBX6 is essential for both PSM progression and IM maturation. Loss of TBX6 arrests PSM at the progenitor stage and blocks activation of the renal developmental program required for functional IM formation.

### Multi-Omics trajectory analysis reveals TBX6-dependent regulatory dynamics of PSM-to-IM differentiation

To dissect the dynamic regulatory changes underlying the PSM arrest and IM maturation defect, we performed integrated single-cell trajectory analysis focused on the PSM-to-IM transition. We first isolated PSM and IM cells from both WT and TBX6-KO scRNA-seq datasets. **(Figures S9A-B)**. To directly compare developmental progression, we aligned WT and KO cells to a unified pseudotime axis using integrated trajectory analysis **(Figure 6A)**.

**Figure 6.**
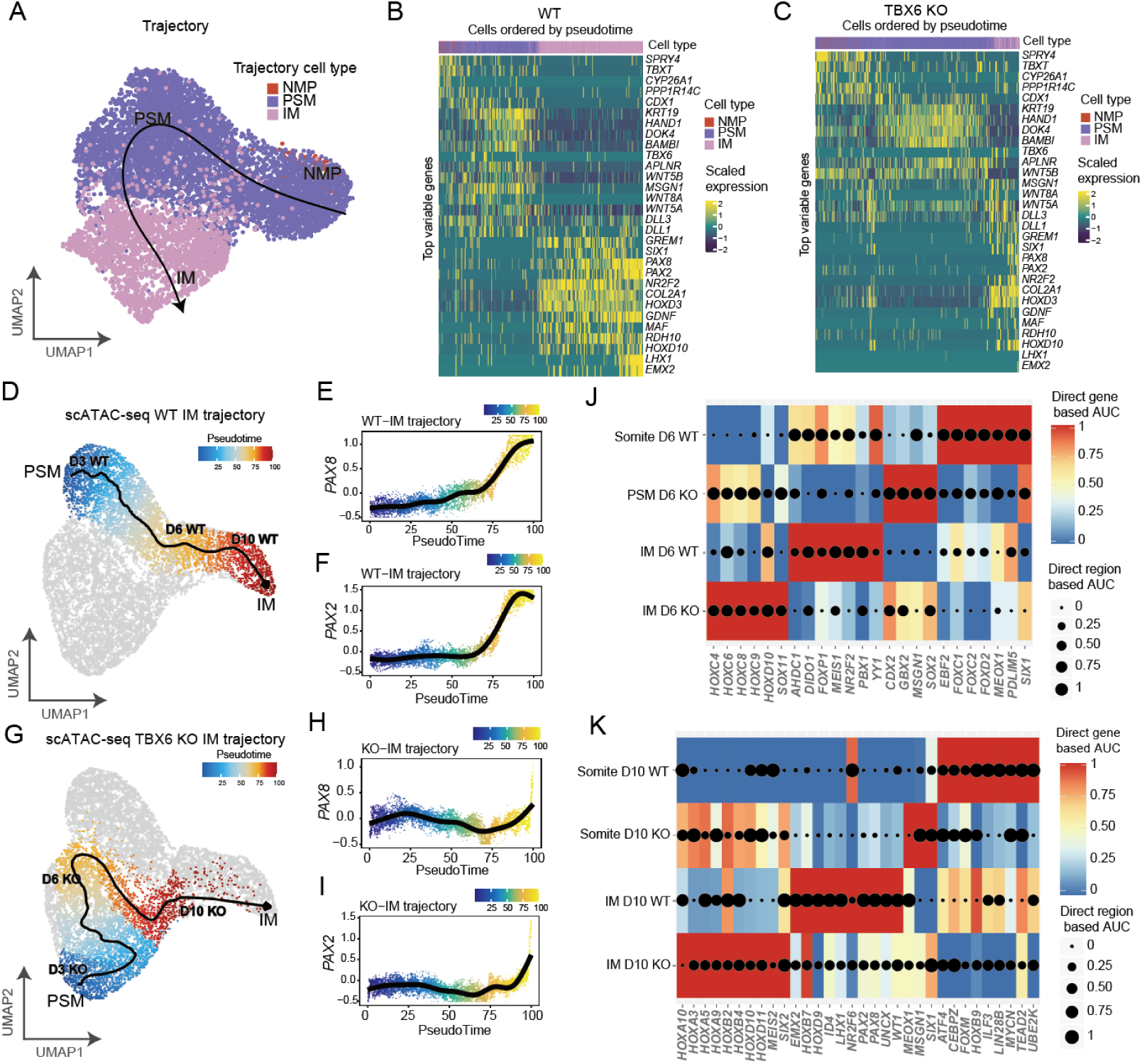
TBX6 disrupts the PSM-to-IM differentiation trajectory and the underlying regulon programs. (A) Slingshot pseudotime trajectory analysis of integrated scRNA-seq data. The trajectory depicts the developmental progression from NMPs/PSM to mature IM cells. (B, C) Pseudotime-ordered heatmaps of top variable gene expression in wild-type (B) and TBX6-KO (C) cells. Rows represent genes, columns represent cells ordered by pseudotime, and colors indicate scaled expression levels. (D) Trajectory analysis of wild-type IM cells from the scATAC-seq dataset. Cells are colored by pseudotime, illustrating the unperturbed progression from PSM (blue) to IM (red) differentiation. (E, F) Chromatin accessibility dynamics of key IM regulators PAX8 (E) and PAX2 (F) along the pseudotime axis in wild-type cells. (G) Trajectory analysis of TBX6-KO IM cells from the scATAC-seq dataset. Cells are colored by pseudotime, illustrating the arrested and disorganized from PSM (blue) to IM (red) differentiation. (H, I) Chromatin accessibility dynamics of key IM regulators PAX8 (H) and PAX2 (I) along the pseudotime axis in TBX6-KO cells. (J, K) Regulon activity analysis comparing direct gene-based and region-based AUC scores of key transcription factor regulons in PSM, IM, and somite cells from NMO day 6 (J) and day 10 (K) wild-type and TBX6-KO conditions.

WT cells spanned the entire differentiation trajectory, and the majority progressed toward mature IM identity. In contrast, TBX6-KO cells were strongly enriched at the PSM progenitor stage **(Figures 6B-C)**. Consistently, gene expression dynamics revealed a clear, ordered program in WT cells, with PSM genes downregulated and IM genes upregulated along the trajectory. In TBX6-KO cells, however, this dynamic program was markedly perturbed: the normal upregulation of IM-specific genes was largely abolished, and PSM signature genes remained persistently high, indicating a failure to exit the progenitor state and activate the downstream differentiation program **(Figures 6B-C)**.

To further explore the chromatin regulatory changes underlying these transcriptional defects, we performed parallel scATAC-seq analysis, focusing again on the PSM and IM lineage **(Figure 6D)**. Pseudotime trajectory analysis on the scATAC-seq data revealed that a smooth, unidirectional trajectory from PSM to IM, with the key IM transcription factors PAX8 and PAX2 exhibiting a progressive increase in their regulatory activity along the pseudotime axis in WT samples **(Figures 6E,F)**. In TBX6-KO samples, the trajectory was markedly disorganized, with abnormal branching **(Figure 6G)**. Most cells were arrested at the early PSM stage, and the normal upregulation of PAX8 and PAX2 activity was largely abolished, with only a small fraction of cells showing activation of these IM regulators, confirming that the chromatin-level activation of the IM program was dependent on TBX6 **(Figures 6H, I)**. The heatmap of full regulatory dynamics further confirmed this pattern: WT cells showed a clear, ordered transition of regulatory activity along the trajectory, while KO cells exhibited a disrupted and stalled regulatory program **(Figures S9D-E)**.

Next, we compared regulon activity in PSM, somite, and IM populations from WT and TBX6-KO NMOs at day 6 and day 10 **(Figures 6J, 6K)**. At day 6, WT NMOs exhibited clear, lineage-restricted regulon programs. In WT IM cells, IM regulators such as PBX1 and NR2F2 were beginning to be activated, reflecting the onset of IM fate commitment. In contrast, TBX6-KO IM cells failed to undergo this early regulatory transition, retaining a stable PSM-like regulon signature with persistent high activity of HOX factors and CDX2, while early IM-specific regulons remained largely inactive. By day 10, the regulatory divergence between WT and KO was further amplified. In WT NMOs, IM cells had fully activated the mature IM regulatory program, with robust activity of key IM transcription factors including PAX2, PAX8, and FOXD1. In TBX6-KO NMOs, IM-specific regulons—including PAX2, PAX8, and FOXD1—remained minimally active, failing to upregulate even at this late stage when WT IM cells are fully committed,consistent with these multi-omic trajectory results.

Taken together, these multi-omics analyses demonstrate that TBX6 is essential for the proper progression of the PSM-to-IM differentiation trajectory. Loss of TBX6 blocks the exit of progenitors from the PSM state, and prevents the step-wise activation of the chromatin and transcriptional regulatory programs that drive IM maturation, providing a mechanistic explanation for the severe and persistent defects in IM and renal development observed in TBX6-deficient organoids. These findings also provide a molecular basis for the clinical observations that TBX6 mutations in human patients are associated with both somite defects and congenital kidney abnormalities^16,17,51^, highlighting the critical role of TBX6 in coordinating the development of these two lineages.

## DISCUSSION

The development of the human posterior trunk relies on neuromesodermal progenitors, but the lineage hierarchy of their mesodermal derivatives and the molecular control of their fate decisions remain poorly defined^2,35^. In this study, we combined single-cell Cas9-Tracer lineage tracing with multi-omic profiling in human neuromesodermal organoids to resolve these processes at clonal resolution. We show that intermediate mesoderm and paraxial mesoderm arise from a shared presomitic mesoderm progenitor downstream of NMPs, while lateral plate mesoderm segregates early from committed mesodermal progenitors downstream of NMPs. We further demonstrate that TBX6 is not only required for somite formation but also essential for the stepwise activation of the IM transcriptional program, providing a mechanistic link between TBX6 mutations and both somite and kidney phenotypes in humans.

Our state-fate mapping confirms that early NMPs are not restricted to neural and paraxial mesoderm fates but can generate LPM *in vitro*. This has critical implications for translational applications: if the goal is to produce posterior LPM derivatives—such as vascular, connective, or body wall tissues—then NMPs represent a viable starting progenitor population. Conversely, for directed differentiation toward IM-derived kidney or PM-derived skeletal lineages, our data imply that culture conditions must actively suppress the LPM fate to avoid off-target lineage contamination. This insight may refine existing protocols and underscores the value of human-specific lineage data for improving the purity of therapeutically relevant cell types.

NMPs are increasingly recognized as a promising source of posterior spinal neurons for treating spinal cord injury or neurodegeneration^52^. However, efficient translation requires precise lineage control. Spinal cord injury and motor neuron disease require desired populations of regionally appropriate posterior neurons with minimal contamination from neural crest or mesodermal lineages^53–55^. Our identification of two distinct neural trajectories from NMP and the early molecular signatures that distinguish them may enable rational enrichment of neuronal progenitors without neural crest drift. Such precision is essential for generating functional, transplantable neuronal populations that match the rostro-caudal identity of the injured spinal cord. By defining the earliest fate choices in human NMP-derived neural development, our study provides a mechanistic foundation for improving the specificity of NMP-based cell-replacement therapies.

Our finding that TBX6 governs both PM and IM development carries significant implications for understanding the pathogenesis of TBX6-associated congenital disorders. In humans, TBX6 variants are linked to both congenital scoliosis (a PM-derived defect) and CAKUT (an IM-derived defect)^21–23^. The dual phenotype has been difficult to reconcile mechanistically, as TBX6 was primarily known as a regulator of paraxial mesoderm specification. Our data now provide a unified explanation: because IM and PM share a common TBX6-dependent PSM progenitor, loss of TBX6 simultaneously disrupts both lineages. This model is further supported by recent work in mouse models demonstrating that the nephric mesenchyme — the embryonic precursor of the kidney — develops from Tbx6-expressing NMP derivatives^18,52^. Our study extends these findings to the human system and adds mechanistic depth by demonstrating that TBX6 is required not only for PSM establishment but also for the regulatory switch that activates the IM transcriptional program. The observation that TBX6-KO IM cells retain PSM-like regulon signatures while failing to activate IM-specific regulators such as PAX2, PAX8, and FOXD1 suggests that TBX6 functions as a gatekeeper of the PSM-to-IM transition, rather than merely as a specifier of paraxial mesoderm identity.

In conclusion, this work provides a high-resolution clonal framework for human posterior mesoderm development, establishing the lineage hierarchy of NMP-derived lineages and uncovering a dual role for TBX6 in PSM progression and IM maturation. By demonstrating that human IM and PM share a common TBX6-dependent progenitor, we provide a mechanistic basis for the combined somite and kidney phenotypes associated with TBX6 mutations. These findings advance our fundamental understanding of human trunk development and highlight the utility of NMOs and lineage tracing for dissecting the origins of congenital disorders. Future work building on this platform will enable further dissection of the signals and transcriptional networks that control mesodermal fate decisions, with implications for disease modeling and regenerative medicine.

### Limitations of the study

Several limitations should be considered. First, while our NMO system recapitulates key features of human posterior neuromesodermal development and shows high transcriptional correlation with published human embryonic data^37^, it remains an *in vitro* model that cannot fully replicate the spatial signaling gradients and tissue–tissue interactions of the native embryonic environment. Second, although our time course extends to day 50, we have not assessed the functional maturation of terminal cell types — for example, whether NMO-derived IM cells can form nephron structures or whether NMO-derived neurons exhibit functional electrophysiological properties. Third, the chromatin accessibility changes and regulatory network perturbations we identify in TBX6-KO cells require direct validation through ChIP-seq or CUT&RUN to establish whether TBX6 directly occupies the enhancers of IM-specific genes, or whether its effect is predominantly indirect through maintaining the PSM chromatin landscape. Finally, our study identified several novel regulators of lineage specification, including the new candidate genes for NC fate decision, but the *in vivo* functional roles of these gene require further experimental validation determined.

## ACKNOWLEDGMENTS

We are grateful to Dr. Baoyang Hu for sharing the H9-iCas9 cell line, Dr. Hui Yang for sharing the piggybac-Cas9-EGFP plasmid, and Dr. Michelle Chan for providing the Cas9 gRNA sequences. We also thank MGI for providing partial free single-cell RNA-seq reagent kits that supported this study. This study was supported by grants from Guangdong Major Project of Basic Research (2026B0303000013), the National Key R&D Program of China (2024YFC3405600), National Natural Science Foundation of China (32270854), Guangdong Basic and Applied Basic Research Foundation (2024B1515040020), Science and Technology Planning Project of Guangdong Province (2023B1212060050, 2023B1212120009), Basic Research Project of Guangzhou Institutes of Biomedicine and Health, Chinese Academy of Sciences (GIBHBRP23-01, GIBHBRP24-01), Major Project of Guangzhou National Laboratory (GZNL2023A03005), Guangdong Provincial Key Laboratory of Stem Cell and Regenerative Medicine (KLRB202303). C.C. was partly supported by National Natural Science Foundation of China (U21A20203 and 32300679), Guangdong Basic and Applied Basic Research Foundation (2022A1515110942), and Science and Technology Program of Guangzhou (2024A04J4274).

## AUTHOR CONTRIBUTIONS

G.D.P. conceived and supervised the entire study. G.D.P., Y.X.L., and C.C. designed the experiments. Y.X.L. conducted the experiments. C.C. performed the bioinformatic analysis. M.S.L established and characterized wild-type NMO culture systems, L.W., C.W.W., Y.Y., D.Y.W contributed to the work. Y.X.L., C.C., and G.D.P. wrote the manuscript.

## DECLARATION OF INTERESTS

The authors declare no competing interests.

### Materials availability

Plasmids and cell lines generated in this study are available from the lead contact upon request.

### Data availability

All sequencing datasets generated in this study are deposited under BioProject accession number PRJCA064621 in national genomic data center. Processed single cell and lineage data are available upon email request.

### Code availability

The instructions and codes to generate figures in this paper can be found at (https://github.com/gpenglab/Cas9Tracer). Additional codes required to reanalyze the data reported in this work is available from the lead contact upon request.

## METHODS

### Cell Lines

hPSC lines (H9) were maintained on Matrigel-coated plates in mTeSR1 medium (Stem Cell Technologies), with medium changed daily. Cells were passaged every 3-4 days using ReLeSR (Stem Cell Technologies) according to the manufacturer’s instructions. HEK293T cells were cultured in DMEM basic media supplemented with 10% FBS. All the cell lines were cultured in humidified incubators at 37℃ in 5% (v/v) CO_2_.

### Cas9-Tracer plasmid design and cloning

The Cas9-Tracer system consists of three core components:(1) iCas9 plasmid: A piggybac plasmid carrying a doxycycline-inducible Cas9 expression cassette, with Cas9 fused to EGFP to enable expression monitoring. (2) sgRNA-Target plasmid: A second PB vector carrying an EF1α promoter-driven mCherry reporter, followed by three Cas9 target sequences in the 3’UTR of the reporter gene. To enable tunable editing dynamics, we generated two variants of this vector: one with medium-efficiency mismatched targets (“333”) and one with high-efficiency perfect targets (“PPP”). We also incorporated a UCOE (ubiquitous chromatin opening element) upstream of the EF1α promoter to prevent transgene silencing during long-term culture. (3) The lentiviral target plasmid was derived from the pSin-EF1α-Flag vector (a gift from Dr. Hongjie Yao) with the following modification: The flag sequence was replaced by the mCherry-Target sequence using Gibson cloning, and the UCOE (ubiquitous chromatin opening element) sequence (synthesized by Sangon Biotech) was inserted before the EF1α promoter to reduce the gene silencing.

Barcode plasmid adding: NheI and BamHI sites were placed before the target region to facilitate the addition of integration barcodes. IntBCs were inserted into the plasmids using the NEBuilder HiFi Assembly Master Mix (NEB).

### Lentiviral packaging

All lentiviruses were produced in HEK293T cells using the second-generation lentiviral system. Before plasmid transfection, 1x 10^7^ HEK293T cells were plated in a 100 mm dish. Plasmid transfection was performed when the cells reached 70-90% confluency. For transfection, 6 μg Target plasmids, 4 μg PSPAX2, and 2 μg PMD.2G plasmids were used plus with 48 μl PEI. After 12 hours, the transfection reagent was removed by washing the plate with a fresh medium. After 48 hours, the lentiviral suspension was collected and filtered through a 0.45 μm filter to remove cell debris.

### Generation of Cas9-Tracer hES cell lines

We infected the AAVS1-iCas9-KI modified cells with lentivirus carrying the integrative barcodes (intBC) library at a high multiplicity of infection (MOI), to introduce heritable, pre-integrated barcodes into each cell. Following infection, we performed fluorescence-activated cell sorting (FACS) to isolate mCherry-positive cells, which carried the complete Cas9-Tracer cassette.

### NMO Differentiation

NMO differentiation was performed as previously described with minor modifications. Briefly, hPSCs were first differentiated into trunk progenitors via 2D culture in basal medium supplemented with FGF2 (10 ng/ml) and CHIR99021 (3 μM) for 3 days. At the end of 2D induction, cells were dissociated using Accutase to obtain a single-cell suspension. On NMO Day 0, 100 μL NMP cell suspension was plated on an ultra-low binding 96-well plate in NMO medium (N2B27 medium with 10 ng/mL FGF2, 3 μM CHIR, 2 ng/mL IGF1, 2 ng/mL HGF) at a density of 8-9×10^4^ cells/ml. After two days of culture, half of the medium was removed and replaced with 100 μL of NMO medium. On NMO Day 4, the organoids were cultured in N2B27 medium, with the medium being refreshed every two days.

For lineage tracing induction, doxycycline (1 μg/ml) was added to the medium at specified time points (e.g., days 4-8 for the PPP line) to induce Cas9 expression and barcode editing. Organoids were cultured for up to 50 days, with medium changed every 2-3 days.

### Immunofluorescence Staining

For adherent NMPs on coverslips: cells were fixed in 4% PFA for 20–30 min, washed three times with PBS, permeabilized in PBS + 0.3% Triton X-100 at 4 °C for 15 min, and blocked with 5% BSA-containing PBST for 1 h at room temperature. Samples were incubated with primary antibodies overnight at 4 °C, rinsed three times in PBST, then incubated with fluorophore-conjugated secondary antibodies for 1–2 h in dark. After final PBST washes, coverslips were mounted with DAPI-supplemented Fluoroshield medium (Abcam). For NMOs: organoids were fixed in 4% PFA (30 min–3 h, size-dependent), PBS-washed, and dehydrated overnight in 30% sucrose at 4 °C. Organoids were embedded in OCT, frozen at −80 °C, and cut into 10 μm sections using a Leica CM1950 cryostat. Sections were permeabilized and blocked following identical conditions as NMP samples, subjected to sequential primary and secondary antibody incubation, PBST washing, DAPI counterstaining and mounting.

### TBX6 KO Cell Line Generation

We designed two sgRNAs targeting the third and fourth exons of the human TBX6 gene (sgRNA1: 5’-CTGGATTCGTTCCTCTCCG-3’, sgRNA2: 5’-CCAGGGCCCGCTACTTGTT-3’). For transfection, H9-iCas9-KI hESCs were harvested as single cells, and 1 million cells were transfected with the TBX6-KO sgRNA plasmid. Transfection was performed using the Lipofectamine Stem cell transfection reagent (Thermo Fisher), following the manufacturer’s optimized protocol for stem cells. Briefly, 9μg plasmids were mixed with the Lipofectamine Stem Cell reagent in Opti-MEM medium, incubated for 10 minutes at room temperature, and then added to the cell culture medium. After transfection, the medium was replaced with fresh mTeSR1 medium after 24 hours to remove the transfection reagent. After 4 days culture, individual clones were picked into 48-well plates for expansion. Genomic DNA was extracted from each clone. Genotyping was performed by PCR amplification using the forward primer F1 and reverse primer R1, which flank the targeted deletion region. Positive homozygous KO clones were further validated by Sanger sequencing of the PCR products.

### Single-cell RNA-seq

For single-cell preparation, NMOs were dissociated by the Papain dissociation kit (Worthington). In brief, NMOs were incubated in 1 ml Enzyme A along with 20 μl DnaseI at 37℃ for 20-25 min, followed by gentle blowing every 5 minutes with wide-bore tips. After washing and filtering, a suitable volume of PBS with 0.04% BSA was added to resuspend the cells to achieve a cell concentration of 2×10^6^ /ml. Single-cell RNA libraries were prepared using DNBelab C4 kit (BGI-MGI) according to the user’s guide. The main steps of library preparation include droplet formation, demulsification, reverse transcription, second-strand synthesis, cDNA, and oligo product amplification following size selection. Each obtained cDNA pool was divided into two parts. One part of cDNA was prepared by fragmentation, end repair, adaptor ligation, and PCR to get a cDNA library for scRNA-seq. Another part of cDNA was used for amplicon sequencing. Corresponding oligo libraries were generated by PCR amplification from oligo products. The cDNA libraries and oligo libraries were sequenced on the MGI-seq T7 platform.

### Single-cell ATAC-seq

For single-cell ATAC-seq preparation, NMOs were first dissociated into single cells using Accutase (Stem Cell Technologies) at 37°C for 7-10 minutes, with gentle pipetting every 3 minutes to ensure complete dissociation of the organoids. After dissociation, cells were washed with cold PBS, and then subjected to nuclei extraction following the optimized 10x Genomics nuclei isolation protocol. Briefly, cells were lysed in cold lysis buffer (10 mM Tris-HCl pH 7.4, 10 mM NaCl, 3 mM MgCl2, 0.1% NP40, 1 mM DTT, 1 U/µl RNase inhibitor) for 5 minutes on ice, followed by washing and filtering to remove cell debris. After nuclei extraction, the nuclei were resuspended in cold Nuclei Buffer (PBS supplemented with 1% BSA) which was suitable for downstream library preparation.

Single-cell ATAC-seq libraries were prepared using the DNBelab C series High-throughput Single Cell ATAC Library Preparation Kit (MGI) according to the user’s guide. The main steps of library preparation include chromatin transposition by Tn5 transposase, droplet formation, cell barcoding, library amplification, and size selection, following the manufacturer’s protocol. The qualified ATAC-seq libraries were sequenced on the MGI-seq T7 platform, following the standard paired-end sequencing parameters for single-cell ATAC assays.

### Single-cell amplicon sequencing

We performed amplicon sequencing to capture the Cas9-Tracer target region to obtain both the intBC and CRISPR indel information. For the single-cell cDNA template, a pair of primers (C4-F1:5’-ACACTCTTTCCCTACACGACGCTCTTCCGATCTCGTAGCCATGTCGTTCTGCG-3’, C4-R1:5’-TCTCGTGGGCTCG GAGATGTGTATAAGAGACAGTGCAGGAGCGGATTGCTTCGAACC-3’) suitable for the MGI C4 cDNA library adaptors were used for amplification. 50-100 ng cDNA templates were used per amplicon reaction, and two parallel reactions were performed for one sample to reduce potential PCR biases. PCR was performed using 2x Kapa HIFI Mix according to the following program: 95℃, 3 min; 98℃, 15 s; 60℃, 18 s; 72℃, 15s. The first round of PCR was 8-12 cycles and the second round of PCR was 5-6 cycles. PCR products were selected and purified by AMPure XP beads (Beckman). The amplicon libraries were sequenced on the Illumina Nova PE250 platform or MGI-PE300 platform.

### Single-cell RNA-seq data analysis

For each individual sample, a gene expression matrix was generated using the DNBelab C4 analysis pipeline. The Seurat package was used for further processing and downstream analysis. Briefly, we filtered low-quality cells by inspecting the distributions of the number of counts, number of genes, and mitochondrial gene percentage; then we normalized the gene expression matrix, identified the top variable genes, and scaled the data while regressing out confounding factors including cell cycle score, mitochondrial percentage, and number of features. We then used DoubletFinder to remove potential doublets. After performing PCA dimension reduction and cell clustering, we further removed low-quality spurious clusters. For cross-sample analysis, state-fate samples were directly merged, and lineage-tracing samples were merged using SCTransform to remove batch effects. TBX6-KO samples and their corresponding wildtype samples at the same stage were integrated using Harmony. We manually annotated cell types according to canonical markers for wildtype samples and further validated their correlation scores with their counterparts in human embryo datasets. TBX6-KO samples were annotated using the Seurat label-transfer strategy, whereas several cell types were manually adjusted according to gene expression markers and developmental stages. For trajectory analysis, we ran scVelo on individual samples and Monocle3 on the merged samples.

### Single-cell ATAC-seq data analysis

Fragment files were generated using the DNBelab C4 analysis pipeline and then further processed using the ArchR package. In brief, we filtered out low-quality cells with low fragment numbers and low transcription start site enrichment scores. After dropping doublet cells, we performed dimension reduction using iterative latent semantic indexing and visualized the cell distribution using UMAP projection. We used the label-transfer strategy to annotate cell types based on the gene score matrix and the corresponding scRNA-seq gene expression matrix. For better comparison, paired wildtype and mutant fragment files from the same stage were loaded as a single ArchR project, and all downstream analyses were performed on this combined ArchR project. Trajectory analysis on scATAC-seq datasets was performed using Slingshot after merging relevant cell types. Based on cell annotations from ArchR analysis, we used SCENIC+ to perform regulon analysis. Before running the SCENIC+ pipeline, we called topics and obtained cell-topic-fragment relationships using the pycisTopic package, then we performed motif enrichment in binarized topics and differentially accessible regions among different cell types.

### Calling clones, phylogenetic tree reconstruction, and cell-type clustering

Amplicon sequencing data were processed using a modified Cassiopeia preprocessing pipeline. Briefly, we corrected cell barcode sequences in good-quality reads by comparing them with the cell barcode sequences called from the corresponding scRNA-seq datasets. After resolving the read-UMI pairs, we aligned reads to the target reference sequence. We then extracted intBC sequences and collapsed sequences within a Levenshtein distance of 2 bp using Starcode software. We applied further filtering on the obtained cell-intBC-UMI matrix to remove low-quality UMIs, cellBCs, and potentially conflicting intBCs.

We used a custom Jaccard similarity-based graph clustering method to call clones. We first generated a cell-cell Jaccard score matrix from the obtained cell-intBC-UMI matrix. The Jaccard score was calculated among cells using intBCs. We then called clones using Girvan-Newman graph component analysis. Of note, Jaccard scores less than 0.6 were set to 0 to avoid low-confidence connections. For the lineage tracing samples, we further refined the intBC and cellBC inclusion in the called clones using a Cassiopeia internal function, and then reconstructed phylogenetic trees for each clonal lineage using the Cassiopeia-greedy algorithm. We integrated the lineage tree information with the annotations from single-cell transcriptomics and visualized the combined trees using Matplotlib.

For cell-type clustering, we utilized the fate coupling function from the CoSpar package. In short, we defined a lineage cluster as cells sharing the same intBC-mutation combination, integrated the lineage cluster with cell type annotation, and calculated fate coupling scores among different cell types. We then calculated pairwise distances among different cell types using the correlation metric and performed Ward’s hierarchical clustering on these pairwise distances. The obtained cell type hierarchy was then converted into a tree. To make the tree more robust, we repeated the same analysis 100 times by randomly selecting 80% of the cells each time, simulating the effect of cell dropout. We used the consensus tree as the final tree structure and labeled the major branches of the tree with supporting values.

## SUPPLEMENTARY FIGURE LEGENDS

**Figure S1.**
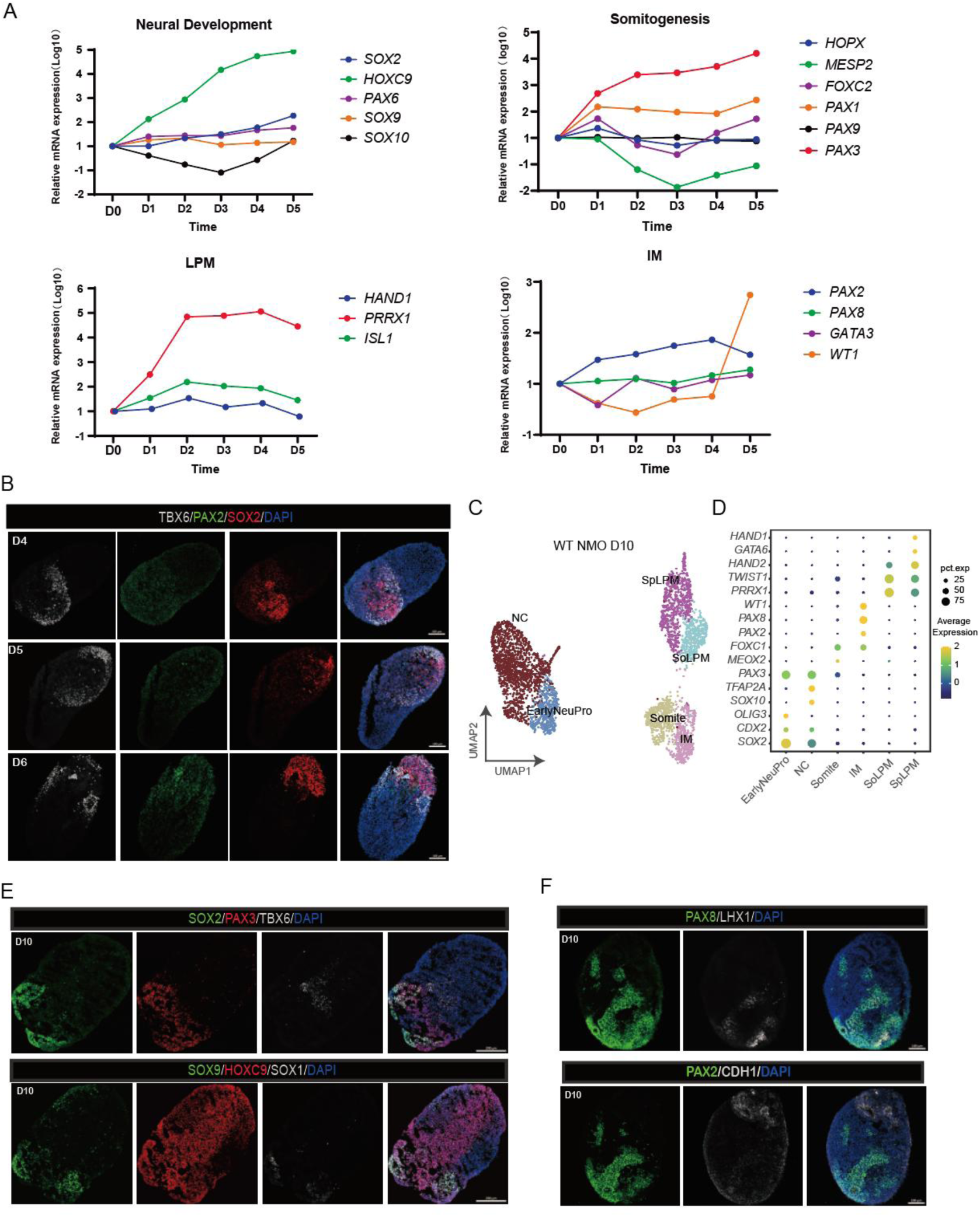
Characterization of wild-type NMO differentiation, related to Figure 1. (A) Time-course qPCR analysis of lineage-specific markers during early NMO differentiation, confirming the dynamic upregulation of canonical markers for neural, paraxial mesoderm, intermediate mesoderm, and lateral plate mesoderm lineages. (B) Immunofluorescence staining of wild-type NMOs collected at D4 (top), D5 (middle), D6 (bottom). Scale bar, 100 ìm. (C) UMAP of scRNA-seq data from day 10 WT NMOs, colored by manually annotated cell types. (D) Dot plot showing the expression of canonical cell-type marker genes in NMO D10. (E, F) Immunofluorescence staining of day 10 wild-type NMOs. Scale bar, 100 μm.

**Figure S2.**
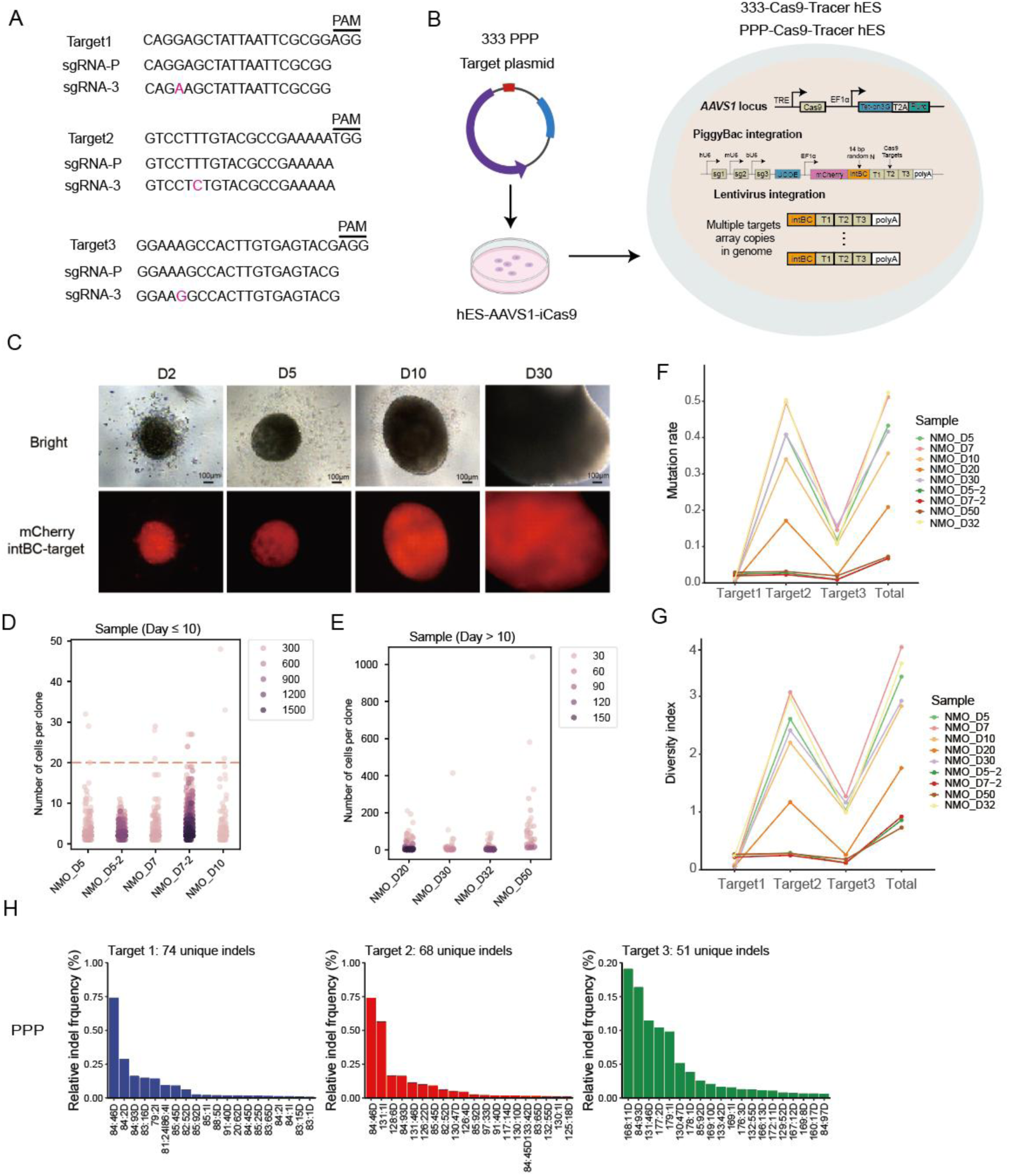
Quality validation of the Cas9-Tracer system for lineage tracing, related to Figure 1. (A) sgRNA target sequences for the 333 and PPP cell lines, with sgRNA-target mismatches highlighted in red. (B) Schematic of the generation of Cas9-Tracer hPSC lines. (C) Brightfield (top) and mCherry (bottom) fluorescence images of 333-Cas9-Tracer NMOs at day 2, 5, 10, and 30 of differentiation, showing stable reporter expression during organoid maturation. Scale bars: 100 μm (D2, D5, D10), 120 μm(D30). (D, E) Scatter plots showing the distribution of cell counts per clone across NMO samples. (D) Early time points (Day ≤ 10) (E) Late time points (Day > 10). (F) Line plot showing the Cas9 editing rate across different target sites, confirming consistent editing efficiency across samples. (G) Line plot showing the barcode diversity index across different target sites for the 333 and PPP cell lines. (H) Bar plot showing the relative frequency of unique indels in the PPP Cas9-Tracer cell lines. Mutations were arranged in descending order of their read fractions, and the enumeration of unique indels was performed when the cumulative fraction first exceeded the 0.95 threshold.

**Figure S3.**
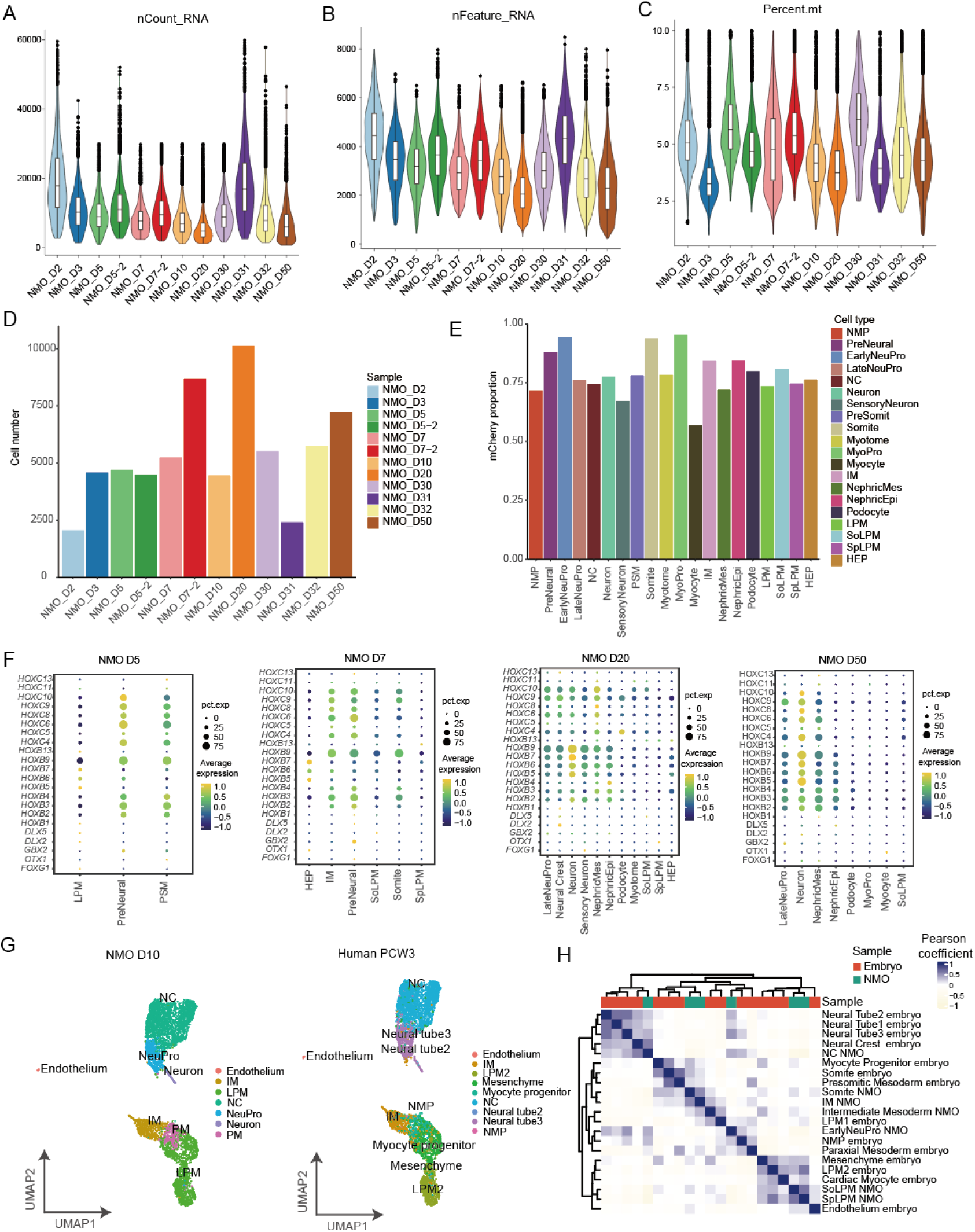
Quality control and in vivo fidelity assessment of the Cas9-Tracer scRNA-seq dataset, related to Figure 1. (A) Violin plot showing the distribution of total RNA counts per cell across all 12 samples, confirming consistent sequencing depth. (B) Violin plot showing the distribution of detected genes per cell across all 12 samples, confirming uniform gene detection rate. (C) Violin plot showing the distribution of mitochondrial read percentage per cell across all 12 samples, confirming high cell quality. (D) Bar plot showing the number of cells recovered from each of the 12 samples after quality filtering. (E) Stacked bar plot showing the proportion of mCherry expressing cells across all cell types. (F) Dot plots showing the expression of canonical anterior-posterior marker genes across cell types at four representative differentiation time points (D5, D7, D20, D50). (G) UMAP of scRNA-seq data from day 10 (D10) NMO (left) and human PCW3 (right), colored by cell type annotations. Cell types in the NMO dataset were mapped to their corresponding *in vivo* human embryonic counterparts. (H) Pearson correlation heatmap comparing the transcriptomes of D10 NMO-derived cell types with primary human PCW3 cell types.

**Figure S4.**
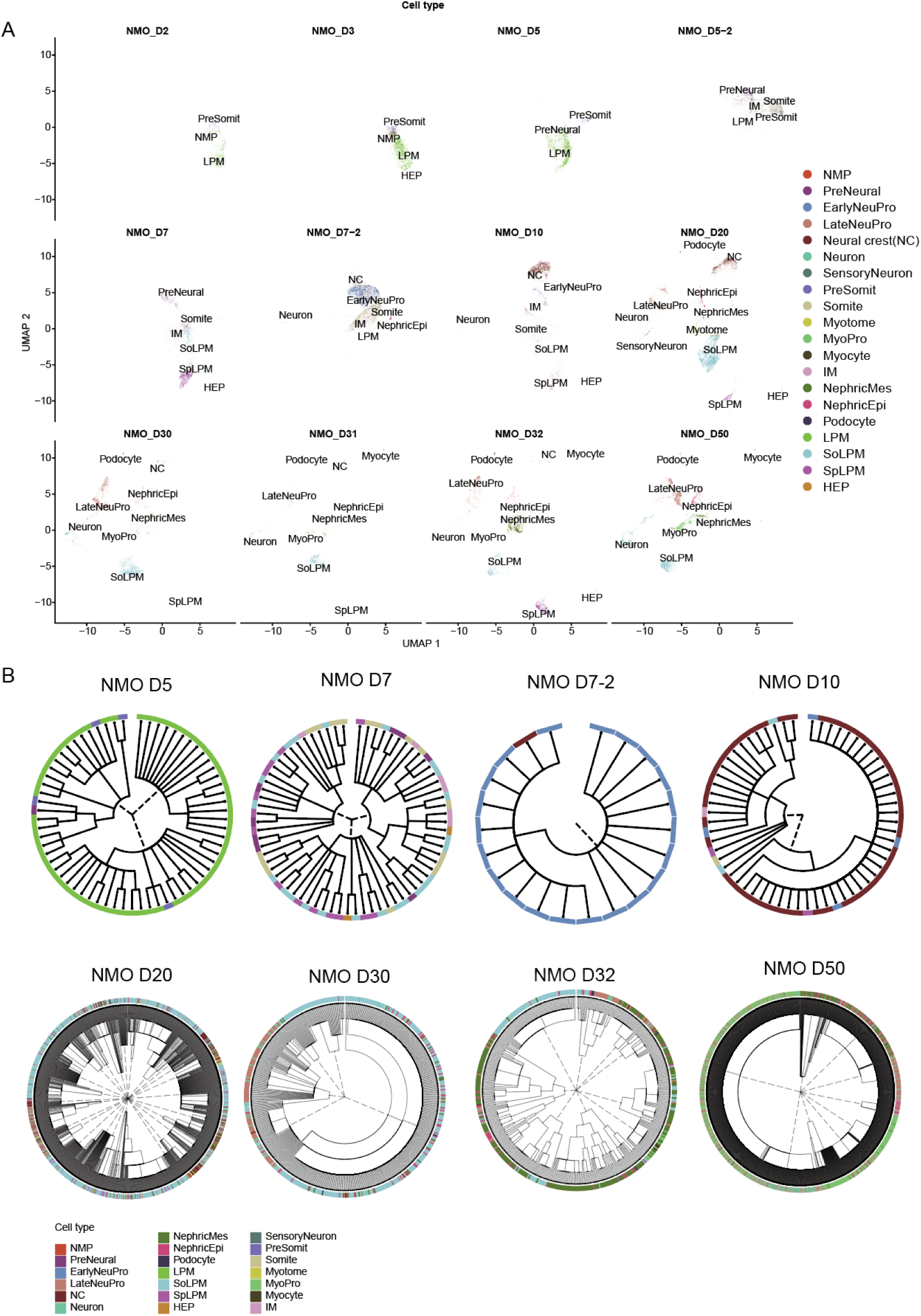
Temporal progression of cellular diversity and clonal dynamics during NMO differentiation, related to Figure 2. (A) UMAP of scRNA-seq data from individual NMO samples collected across differentiation time points (D2 to D50), colored by annotated cell type. (B) Phylogenetic trees constructed from barcode data of representative NMO samples (D5 to D50). Each tree depicts the lineage relationships between cells, with outer rings colored by terminal cell type.

**Figure S5.**
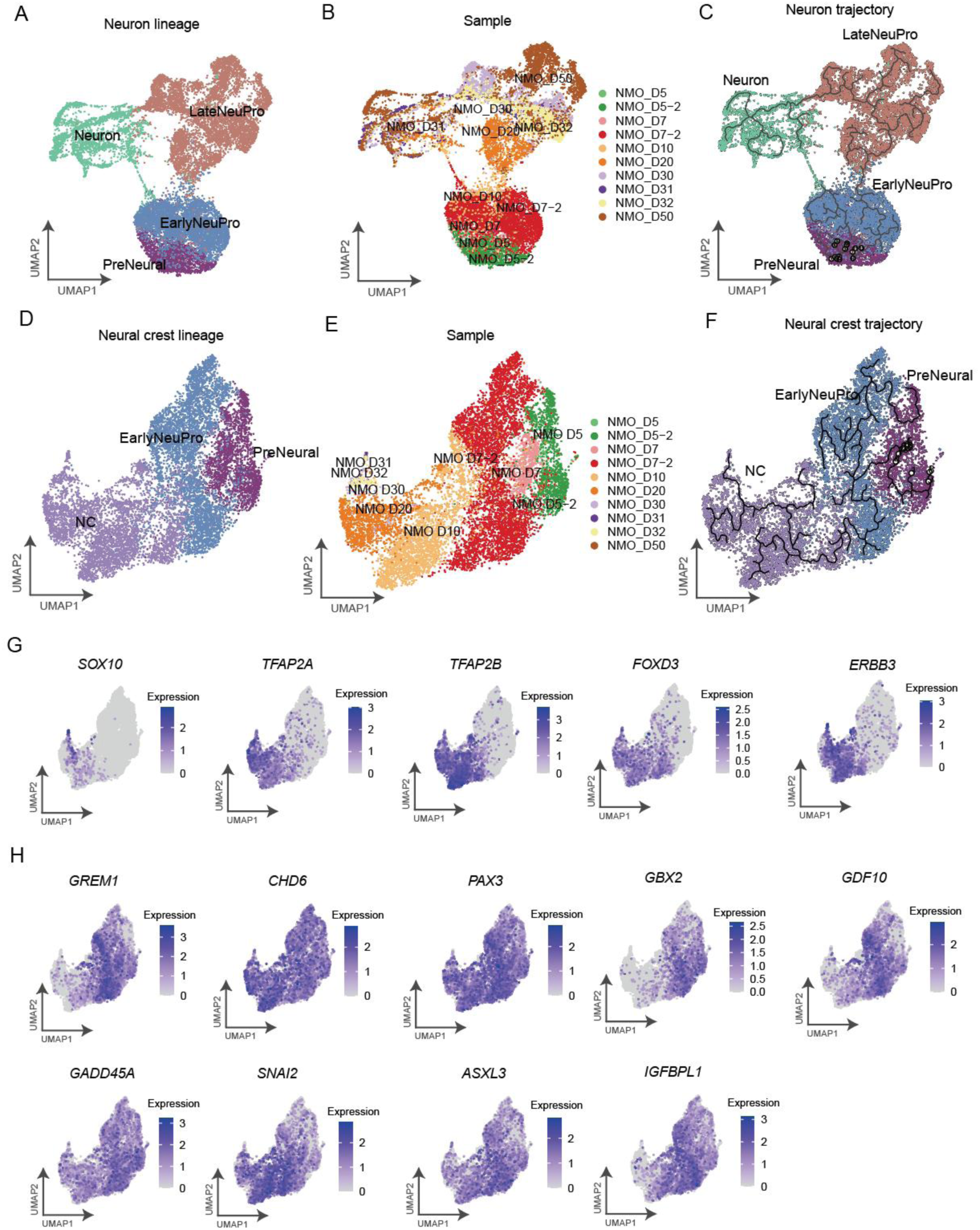
Characterization of neuron and neural crest lineages in NMOs, related to Figure 2. (A) UMAP of the neuron lineage, colored by annotated cell type. (B) UMAP of the neuron lineage, colored by sample/timepoint. (C) The same UMAP as in (A), with inferred differentiation trajectories overlaid, highlighting the path from PreNeural progenitors to mature neurons. (D) UMAP of the neural crest (NC) lineage, colored by cell type. (E) UMAP of the neural crest (NC) lineage, colored by sample/time point. (F) The same UMAP as in (D), with inferred differentiation trajectories overlaid, highlighting the path from PreNeural progenitors to NC cells. (G) Feature plots showing the expression of key neural crest marker genes. (H) Feature plots showing the expression of top DEGs between NC-committed and non-NC-committed EarlyNeuPro clones (related to figure 2L).

**Figure S6.**
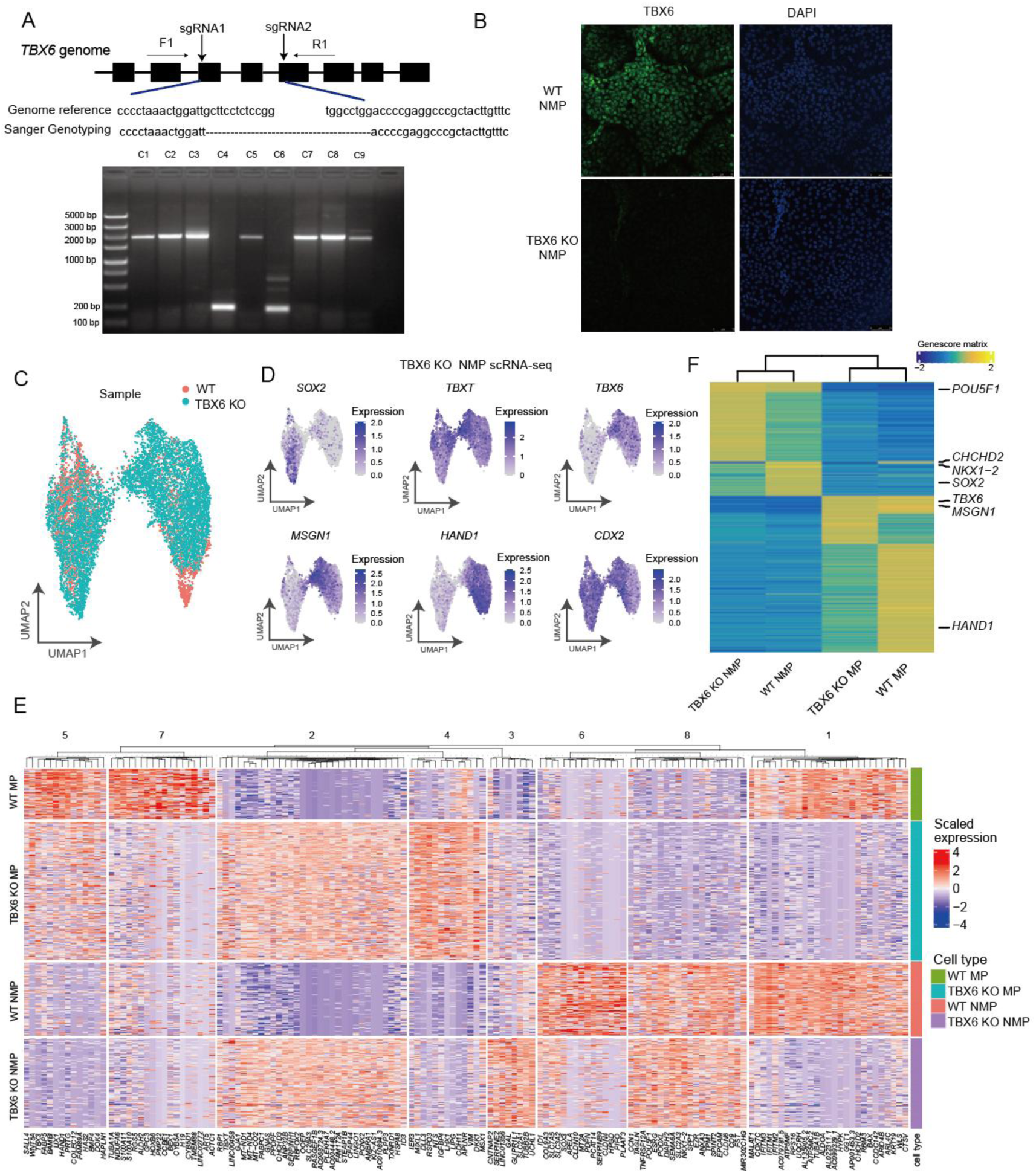
Generation and validation of TBX6-KO NMPs and transcriptomic profiling, related to Figure 5. (A) Schematic of the TBX6 knockout strategy and the genotyping strategy (primers F1, R1). Below, agarose gel electrophoresis of PCR products from clonal cell lines (C1–C9) confirms successful gene editing and deletion of the TBX6 exon2-exon3 region. (B) Immunofluorescence staining of wild-type and TBX6 KO NMP for TBX6 (green) and DAPI (blue), confirming the loss of TBX6 protein expression in KO cells. (C) UMAP of scRNA-seq data from wild-type and TBX6 KO NMPs at ?h, colored by genotype. (D) UMAP feature plots showing the expression of key NMP and mesodermal marker genes (*SOX2, TBXT, TBX6, MSGN1, HAND1, CDX2*) in the TBX6 KO 72 h(hours) NMP scRNA-seq data. (E) Heatmap showing scaled gene expression in wild-type and TBX6 KO 72 h(hours) NMP scRNA-seq data. (F) Heatmap of activity scores for key transcription factors in WT and TBX6 KO 72 h(hours) NMP scATAC-seq data.

**Figure S7.**
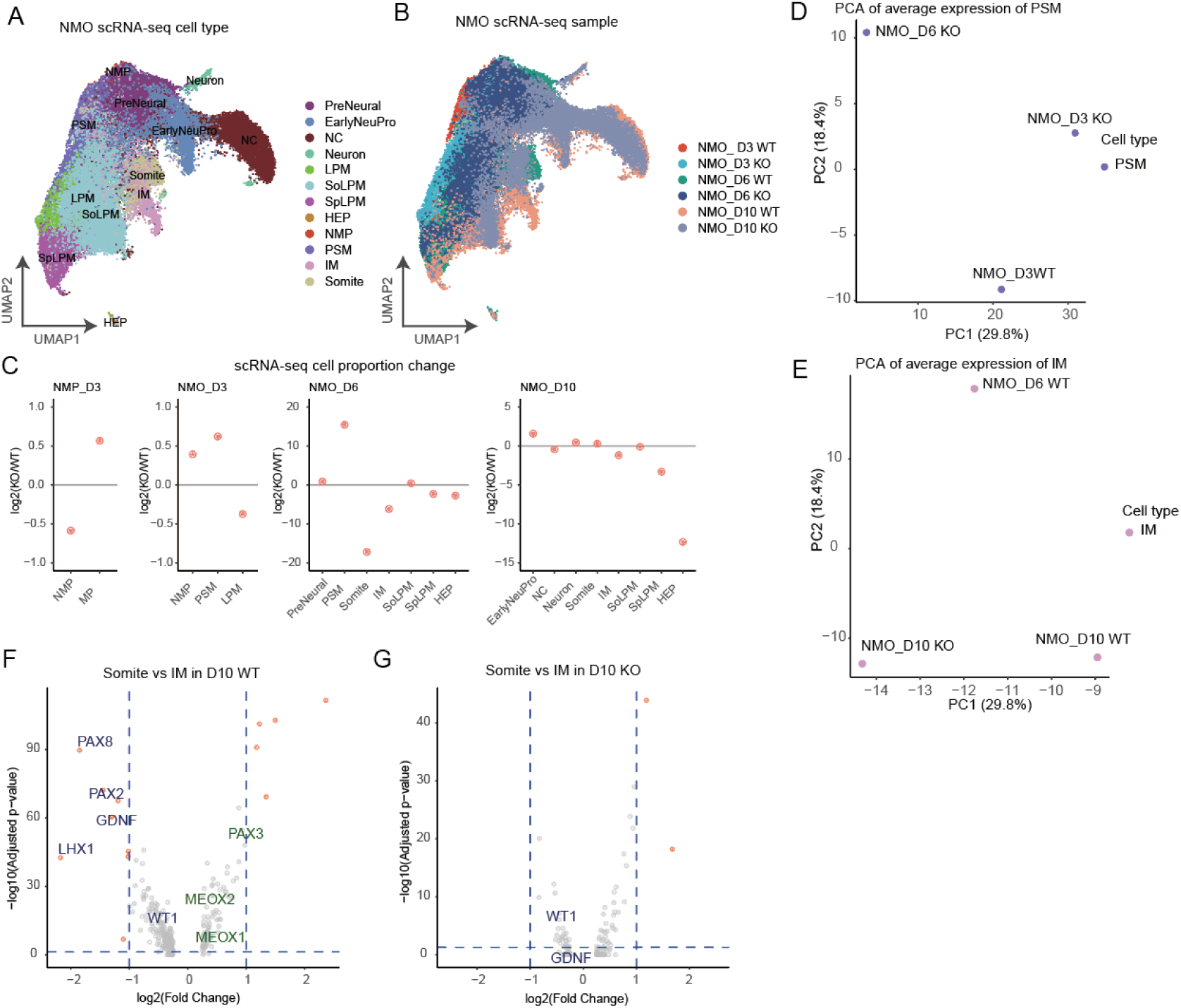
Cellular composition and transcriptomic alterations in TBX6-KO NMOs, related to Figure 5. (A) UMAP visualization of integrated scRNA-seq data from all NMO samples, colored by annotated cell type. (B) UMAP visualization of integrated scRNA-seq data from all NMO samples, colored by sample and genotype (WT and TBX6-KO). (C) Scatter plots showing log_2_ (fold changes) in cell type proportions between wild-type and TBX6-KO NMP and NMOs at day 3, 6, and 10, based on scRNA-seq samples. (D, E) Principal component analysis (PCA) of average gene expression profiles of PSM (D) and IM (E) cells, comparing wild-type and TBX6-KO samples at different time points. (F, G) Volcano plots showing differential gene expression between somite and IM cells in day 10 wild-type (F) and TBX6-KO (G) NMOs. Key IM and somite marker genes are labeled.

**Figure S8.**
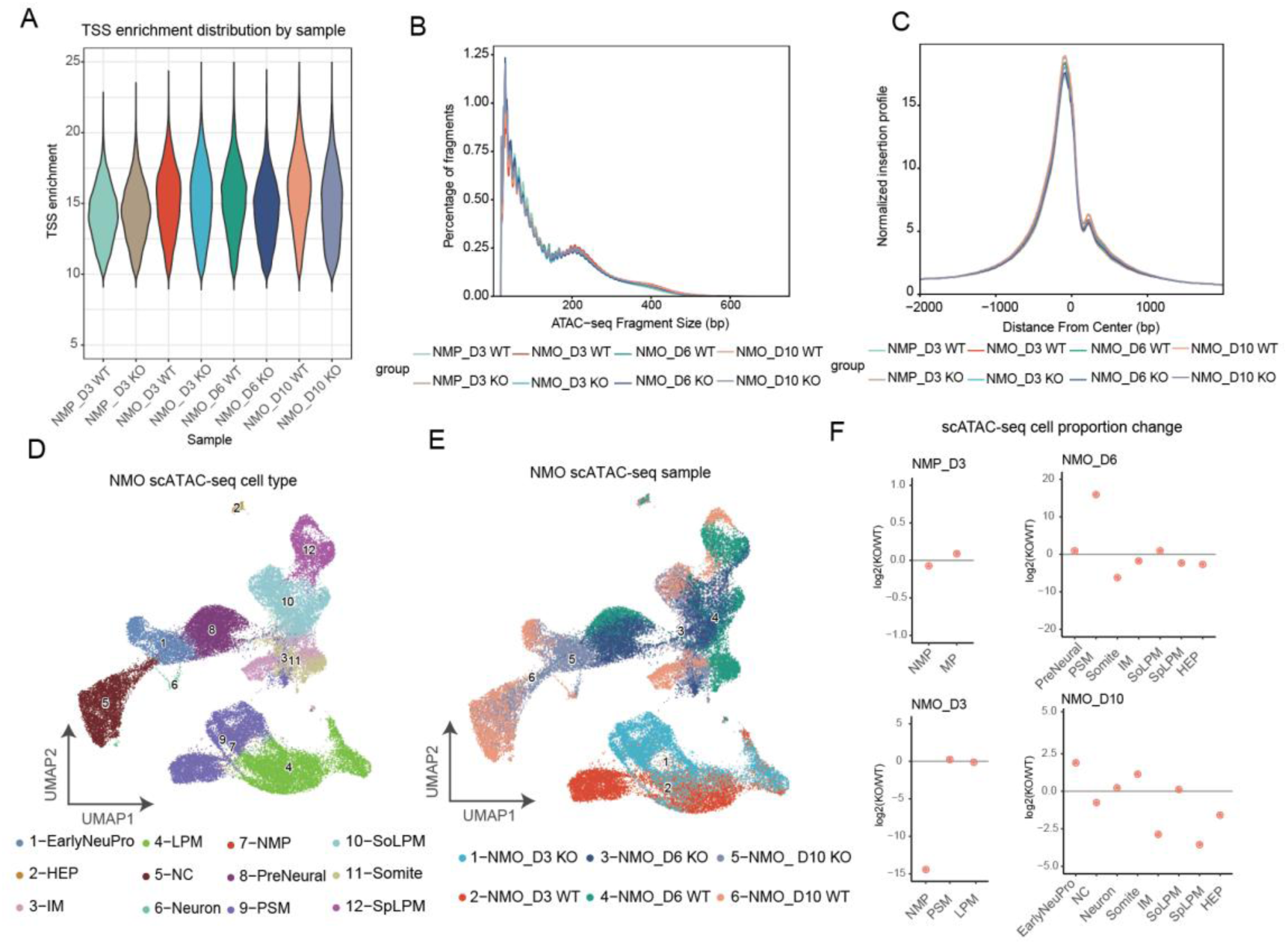
Quality control and profiling of the scATAC-seq dataset in WT and TBX6-KO NMOs, related to Figure 5. (A) Violin plot showing the distribution of TSS enrichment scores across all scATAC-seq samples, confirming consistent chromatin accessibility quality. (B) Fragment size distribution plot showing the percentage of fragments of different lengths across wild-type and TBX6-KO samples, confirming typical nucleosome-free and mono-nucleosomal fragment patterns. (C) Normalized insertion profile centered at transcription start sites (TSS), showing strong enrichment of insertions at the TSS across all samples. (D) UMAP visualization of integrated scATAC-seq data, colored by manually annotated cell type. (E) The same UMAP as in (D), colored by sample and genotype (wild-type vs. TBX6-KO), showing consistent cell type clustering across conditions. (F) Scatter plots showing log_2_ (fold changes) in cell type proportions between wild-type and TBX6-KO NMP and NMO at day 3, 6, and 10, based on scATAC-seq samples.

**Figure S9.**
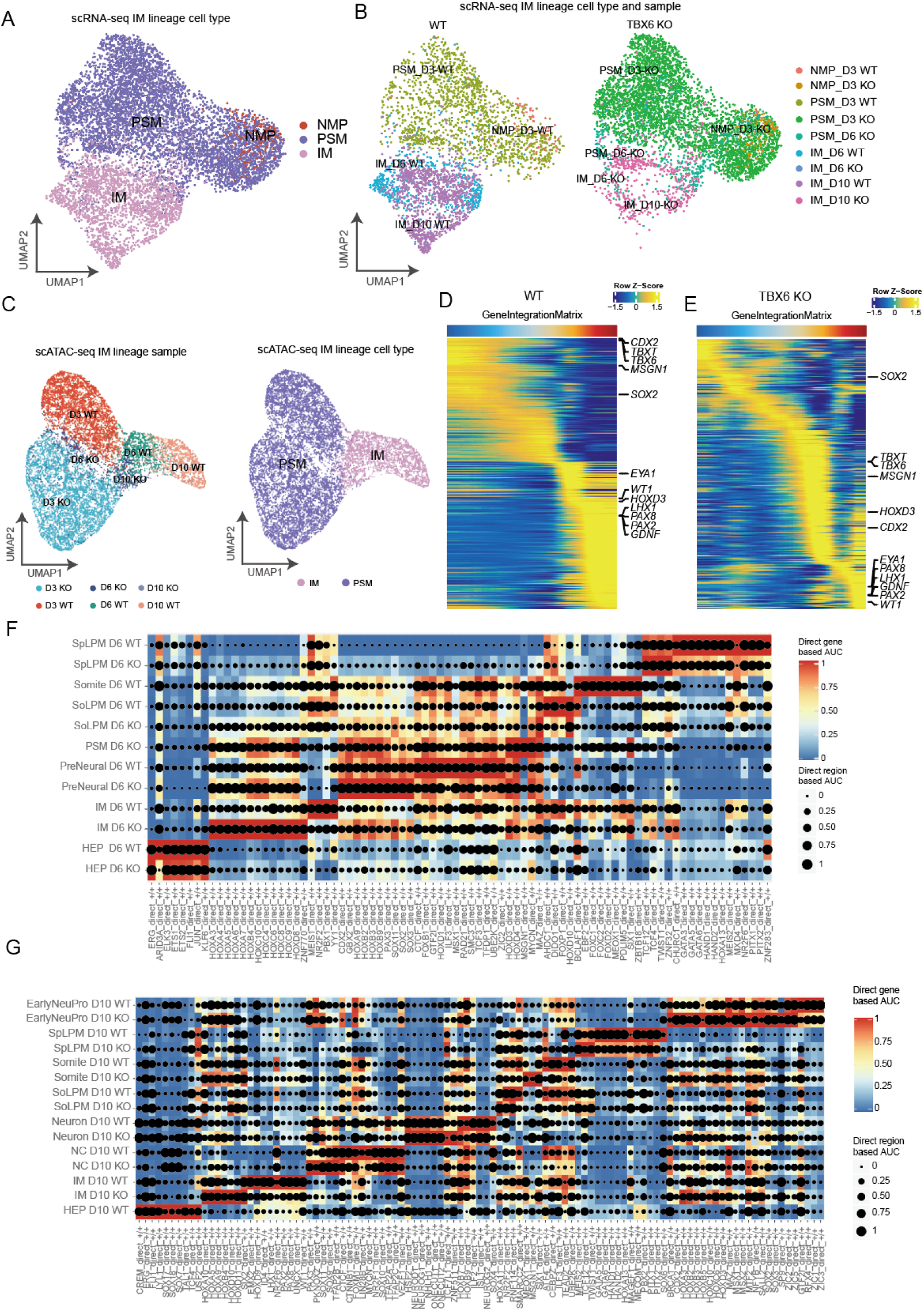
Multi-omics profiling and regulon analysis of the PSM-to-IM trajectory in WT and TBX6-KO NMOs, related to Figure 6. (A) UMAP of scRNA-seq data from the PSM-to-IM lineage, colored by cell type. (B) UMAP of scRNA-seq data from the PSM-to-IM lineage, colored by sample, genotype (wild-type and TBX6-KO), and time point. (C) UMAP of scATAC-seq data from the PSM-to-IM lineage, colored by sample/time point (left) and cell type (right). (D, E) Pseudotime-ordered heatmaps showing scaled gene expression dynamics along the PSM-to-IM trajectory in wild-type (D) and TBX6-KO (E) cells. (F, G) Regulon activity heatmaps comparing direct gene-based and region-based AUC scores of key transcription factors across cell types and sample at day 6 (F) and day 10 (G).

